# Multi-transmitter *high-fidelity* innervation of the forebrain extended amygdala by pontine PACAP-expressing neurons

**DOI:** 10.1101/2025.04.03.647119

**Authors:** Limei Zhang, Vito S. Hernández, David M. Giraldo, Sunny Z. Jiang, Juan C. León-Contreras, Rong Ye, Shiliang Zhang, Martin K-H Schäfer, Rafael A. Barrio, Rogelio Hernández-Pando, Sabina Schoenherr, Francesco Ferraguti, Lee E. Eiden

**Author notes:** Sabbatical researcher (Limei Zhang, Vito S. Hernández). **Author Contributions:** LZ, VSH, FF and LEE conceived and designed the study. LZ, VSH, DMG, SZJ, JCL, RY, SZ, MK-HS, SS and FF performed experiments and collected data. DMG optimized 3D-FIBSEM reconstruction methods. LZ, VSH, RB, FF, LEE analyzed and interpreted the data. LZ, RH-P, RAB, FF, LEE supervised the project and secured funding. LZ wrote the original draft of the manuscript. FF, VSH, DMG, RY, LEE contributed to review & editing. All authors have read and approved the final version of the manuscript. **Competing Interest Statement:** The authors declare no competing interest.

## Abstract

We have discovered a highly specialized innervation of the forebrain by pituitary adenylate cyclase activating polypeptide (PACAP) immunohistochemistry originating from the brain stem that uses glutamate, acetylcholine and PACAP and other peptides as neurotransmitters. The parent neurons of the axons are in the Kölliker-Fuse nucleus and their terminals form calyx-like multi-release site synapses in the rodent forebrain extended amygdala similar to the calyx of Held in the auditory brain stem. The latter is a giant, excitatory, cup-like axo-somatic high-fidelity synapse. The PACAP-positive terminals also form enveloping axo-somatic specialization with mixed glutamatergic and cholinergic molecular identities, co-expressing VGluT1, VGluT2, VAChT, and the neuropeptides PACAP, CGRP, and neurotensin, together with calretinin in the presynaptic compartment. We identified a distinct neuronal subpopulation in the pontine Kölliker-Fuse region of the parabrachial complex that gives rise to these calyceal terminals, which engulf PKCẟ⁺/GluD1⁺ somata in the capsular central amygdala and oval bed nucleus of the stria terminalis. Strikingly, GluD1 immunolabeling is concentrated at axo-somatic contact zones apposed to VAChT⁺ presynaptic vesicle clusters zones but is absent from postsynaptic densities of conventional type I synapses within the same terminals. The results demonstrate a previously unrecognized multimodal calyx-like synapse in the forebrain, with parallel fast ionotropic and modulatory peptidergic neurotransmission mechanisms, which is a substrate for high-fidelity signal transmission within viscerosensory -emotional circuits.

**Significance Statement:** A high-fidelity axo-somatic glutamatergic synapse is exemplified by the calyx of Held in the auditory brainstem, but is unknown in the forebrain. Here we reveal a multimodal calyceal synapse in the extended amygdala formed by glutamatergic afferents of the pontine Kölliker-Fuse nucleus that contain also several neuropeptides. These giant terminals co-package glutamate, acetylcholine, PACAP, CGRP and neurotensin. They envelop select PKCẟ⁺/GluD1⁺ neurons in the capsular central amygdala and oval BNST. These unique forebrain synapses link viscerosensory input from the hindbrain to limbic autonomic centers. Such high-fidelity multimodal transmission is likely to underlie the precise synchronization of emotional and homeostatic control circuits.

## Introduction

Precise and high-fidelity neurotransmission is a hallmark of certain specialized synaptic architectures in the mammalian brain. One of the most striking examples is the calyx of Held, a giant axo-somatic glutamatergic terminal in the auditory brainstem (1), first described over a century ago by Hans Held (2). This unique synapse, formed by globular bushy cells onto principal neurons in the medial nucleus of the trapezoid body (MNTB), is characterized by its expansive presynaptic terminal that engulfs the soma of its postsynaptic target, supporting fast, one-to-one spike transmission with minimal delay and jitter (1, 3, 4). The structure and function of the calyx of Held have provided a foundational model for understanding synaptic reliability, vesicle dynamics, and temporal precision in sensory circuits (5).

The extended amygdala, comprising the capsular central amygdala (CeC) and bed nucleus of the stria terminalis oval (BSTov) nucleus, is a viscerosensory hub for coordinating autonomic and emotional responses (6, 7). These interconnected nuclei receive robust inputs carrying visceral and nociceptive information from the brainstem parabrachial nucleus (PBN) (8–10). While some features of calyx like synapses have been described in hippocampal and cerebellar terminals (11), fully enveloping, axo-somatic synapses have not been reported in forebrain circuits. Similarly, basket-like axosomatic contacts containing calcitonin gene-related peptide (CGRP) (12, 13) (14, 15), pituitary adenylate cyclase activating polypeptide (PACAP) (16), and neurotensin (17) forming type II axo-somatic synapses have been reported (9, 14, 15), but their three-dimensional ultrastructure and presynaptic origin has remained unknown.

Despite the pivotal role of the central amygdala in homeostatic and stress regulation, the organization of its long-range inputs remains poorly understood. Here, using confocal and transmission and tomographic electron microscopy combined with cell-type-specific tracing, we describe a multimodal calyx-like synapse originating from PACAP-rich neurons in the pontine Kölliker–Fuse nucleus, and targeting PKCẟ⁺/GluD1⁺ neurons within the EA, providing the first forebrain example of a calyceal-type terminal. We show that this giant axo-somatic terminal co-packages markers for glutamate, acetylcholine, and multiple neuropeptides. Through such a structurally and neurochemically novel connection the hindbrain can deliver rapid, precisely timed and high-impact viscerosensory signals to limbic autonomic circuits.

## Results

### PACAP-containing ring-like structures identified in the forebrain

To assess the distribution of PACAP containing inputs in the rodent telencephalon, we performed immunohistochemistry and identified a highly selective ring-like innervation pattern in distinct forebrain regions. These perisomatic PACAP-positive structures were especially prominent in the capsular subdivision of the central amygdala (CeC), the oval nucleus of the bed nucleus of stria terminalis (BSTov), and to a lesser extent, scattered within the dorsal and ventral subdivisions of the lateral septal nucleus (see Supplemental information, SI, Fig. 1). Within the amygdaloid complex (Fig. 1, panels As and Bs), we observed heterogeneous PACAP immunoreactivity across multiple amygdaloid nuclei. A dense mesh of fine, varicose PACAP-immunoreactive (PACAP-ir) fibers was distributed in the lateral amygdala (LA, Fig. 1, A and B, B2), postero-dorsal medial amygdala (MePD, Fig. 1, A and B, and B4), posteroventral amygdala (MePV, Fig. 1 B), basolateral (BLA), and basomedial amygdala (BMA, Fig. 1, A and B, B3) amygdala, as well as in the central lateral (CeL) and central medial (CeM) subdivisions of the amygdala (Fig. 1, A and B, B1) forming diffuse *en passant* type innervation patterns. In contrast, the CeC displayed a population of thicker, non-varicose PACAP^+^ fibers forming distinct ring-like axo-somatic arrangements around a subpopulation of neurons (Fig. 1, A, A1, B, and B1, arrows).

**Figure 1.**
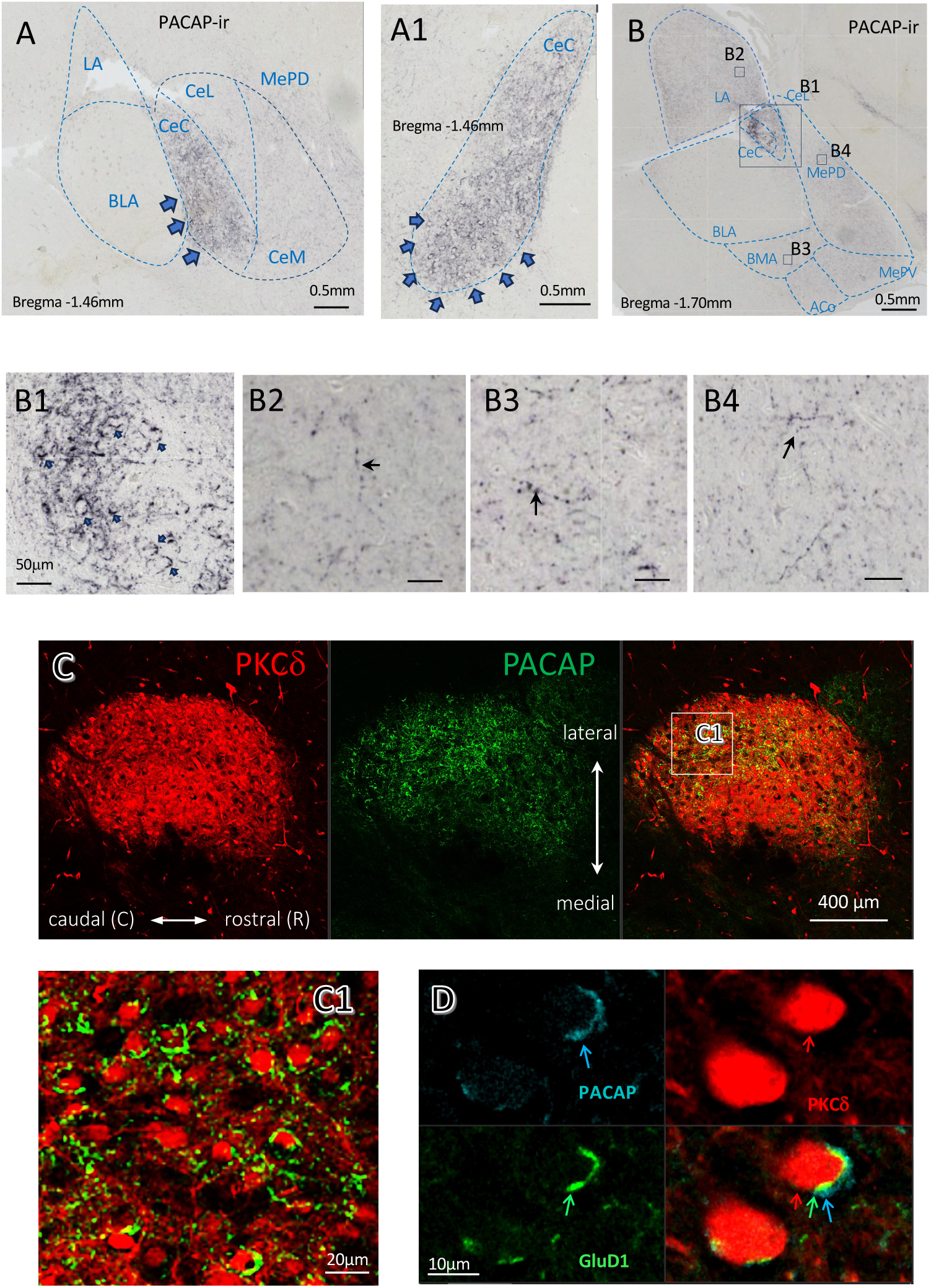
PACAP immunopositive fibers form perisomatic ring-like innervation patterns targeting PKCẟ⁺ neurons in the capsular division of the central amygdala (CeC). **(A–B)** Coronal sections of the amygdaloid complex immunostained for PACAP (PACAP-ir). Blue dashed outlines indicate subdivisions of the amygdaloid nuclei, including central capsular (CeC), central lateral (CeL), central medial (CeM), basolateral (BLA), basomedial (BMA), lateral (LA), and posterodorsal/ posteroventral medial amygdala (MePD/MePV). **(A, A1)** Contralateral amygdaloid complex at approximately bregma - 1.46 mm. Blue-black arrowheads highlight ring-like PACAP-ir structures predominantly in the ventral portion of the CeC. **(B)** Amygdaloid complex at bregma -1.70 mm, where ring-like PACAP+ structures are observed exclusively in the CeC. (B1) Higher magnification of the boxed CeC region in panel B. **(B2 - B4)** Representative PACAP-ir fibers with varicosities (*en passant* type) indicated by black arrows. **(C)** Horizontal section from a Prkcd-Cre::Ai9 mouse showing tdTomato-labeled PKCd+ neurons (red) and PACAP immunostaining (green) in the CeC. **(C1)** Higher magnification of the caudal CeC reveals abundant PACAP-ir perisomatic rings around PKCẟ⁺ cells, mainly located in the posterolateral portion of this horizontal plane. **(D)** High-resolution immunofluorescence for PACAP (blue), GluD1 (green), and PKCẟ-Ai9-tdTomato (red) highlights PACAP-ir labeling around PKCẟ⁺ somata, with GluD1 signals localized to membrane regions in close apposition to PACAP+ elements.

**Figure 2.**
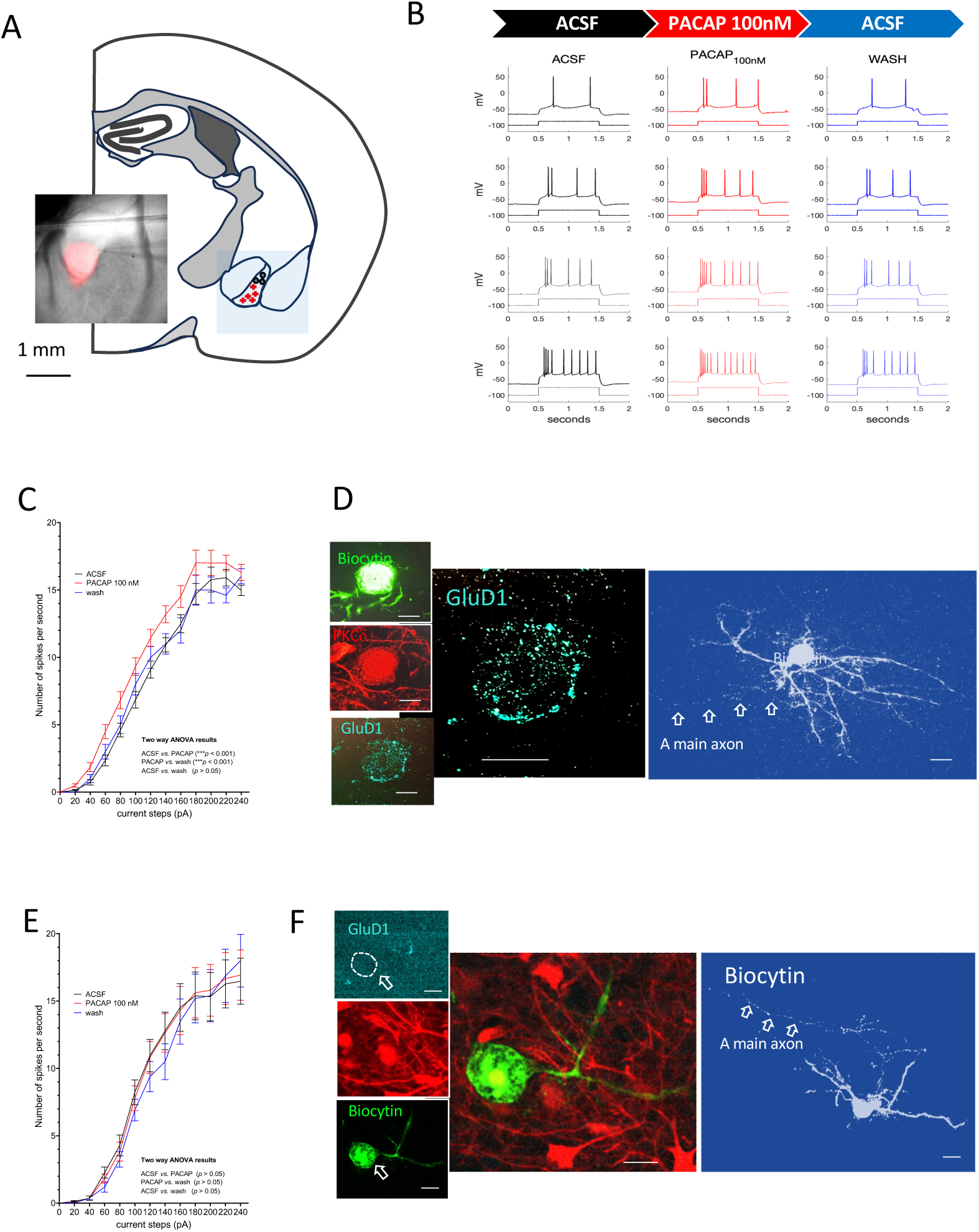
Whole-cell patch-clamp recordings reveal PACAP-sensitive and PACAP-insensitive subtypes of PKCẟ⁺ neurons in mouse central amygdala (CeA). **(A)** Schematic of *ex vivo* recordings performed on horizontal brain slices from Prkcd-Cre::Ai9 mice, in which tdTomato enables identification and targeting of PKCẟ⁺ neurons in the capsular division of the CeA (CeC). Red crosses symbolize the type I PKCẟ⁺ cells, while the grey circles symbolized the type II PKCẟ⁺ (*vide infra*). **(B)** Representative current-clamp traces from a PACAP-sensitive (type I) neuron showing increased excitability during the bath application of 100 nM PACAP (red), which returns to baseline after washout (blue). **(C)** Current-firing frequency (I–F) curves demonstrating enhanced excitability of type I neurons during PACAP application (red curve, n = 21), a two-way ANOVA revealed significant main effects of treatment, F(2,780) = 25.64, p < 0.001. After a 10 min period of washout (blue curve), excitability returned to baseline. **(D)** Example of a type I cell with post hoc immunofluorescence that confirms GluD1 expression (cyan) localized to its plasma membrane (left) and biocytin labeling and its dendritic morphology and axonal projection (right). **(E)** In contrast, PACAP-insensitive (type II) neurons (n=9) showed no change in firing frequency following PACAP application (red curve), no significant effects of treatment were shown in the two-way ANOVA. **(F)** Post hoc immunofluorescence of biocytin filled type II neurons reveals the absence of GluD1 membrane expression (left) despite successful biocytin filling (right). Scale bars: 10 μm.

**Figure 3:**
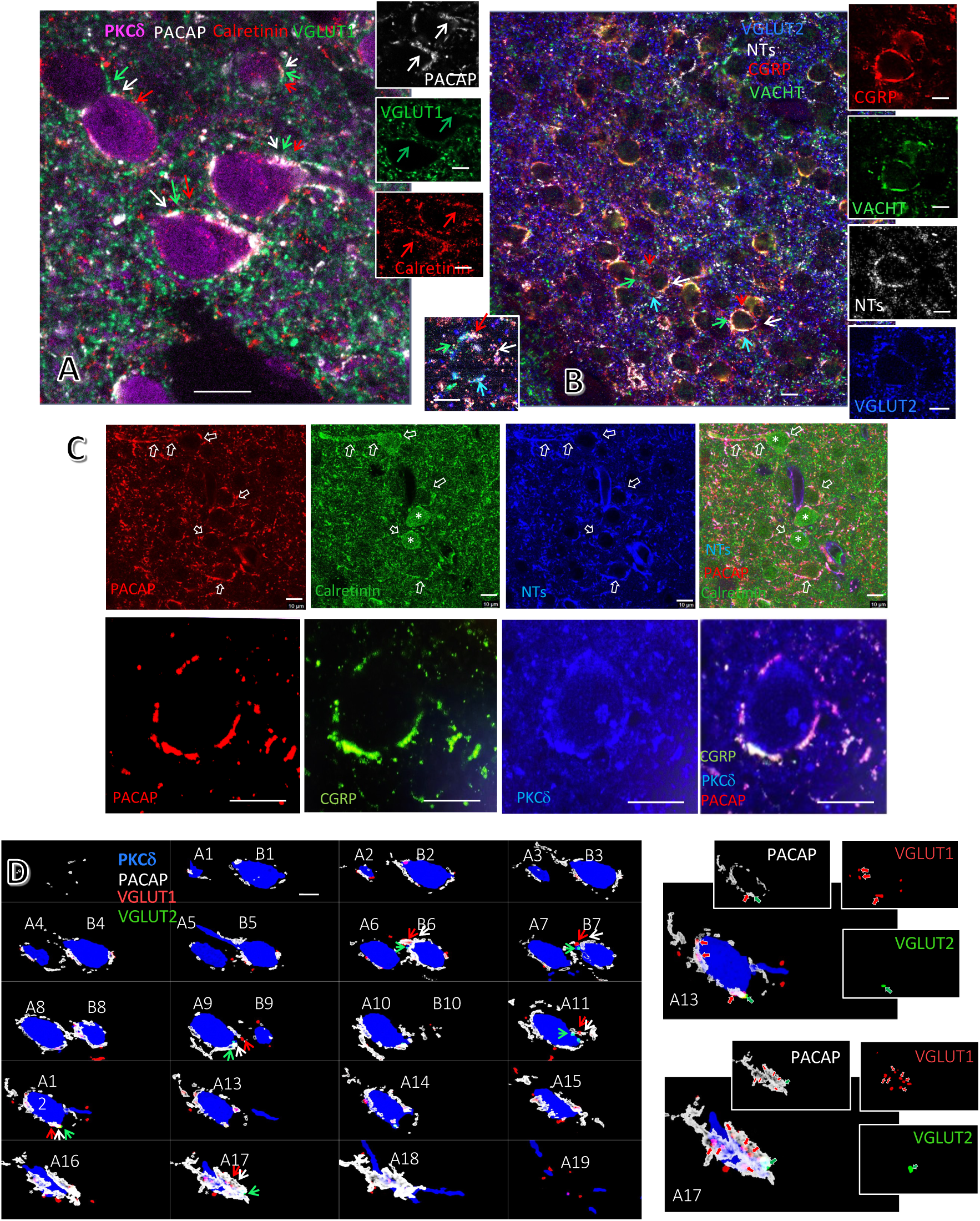
Neurochemical composition of the ring-like structure in rodent CeC. **(A and B):** High-magnification confocal views demonstrate that these perisomatic structures co-express multiple markers, including in A: VGluT1 (green labeling), calretinin (red labeling), PACAP (white labeling) and PKCẟ⁺ (pink labeling); **(B):** neurotensin (NTs, white labeling), vesicular acetylcholine transporter (VAChT, green labeling), VGluT2 (cyan labeling), and calcitonin gene-related peptide (CGRP red labeling). **(C):** top row, co-expression of neuropeptides NTs (blue), PACAP (red) and calretinin (green) forming ring-like structure. Note that some targeted neurons are calretinin expressing (*). In one case, NTs/PACAP/calretinin co-expressed axon approached the targeted cell guided by its main dendrite (arrows at the top part of the panels). Bottom row: ring structure with PACAP/CGRP co-expression engulfing a PKCẟ expressing neuron. **(D)** Segmentation of 20 serial confocal optical sections of two PKCẟ-expressing neurons of rat CeC (blue) illustrates the spatial organization of two VGluT1, VGluT2, PACAP co-expressing ring-like fibers surrounding their somata. Scale bars: 20µm.

### Neuroanatomical characterization of PACAP+ calyceal terminals

To determine the postsynaptic target of these PACAP containing rings, we first assessed somatostatin (SST)-expressing neurons, but these lacked expression of the PAC1 receptor (SI Figure 2, panel B). Since both the CeC and BSTov are known to also contain a high density of PKCẟ-expressing neurons (18, 19), we assessed this population first with RNAscope multiplex method, finding that a subpopulation of *Prkcd* expressing neurons mainly located in the ventro-lateral segment of CeC co-expressed *Adcyap1r1* (PAC1,) *Grid1* (GluD1), *Chrna7* (nAChRa7) (SI Fig. 2, A). Using a *Prkcd*-Cre::Ai9 double transgenic mice, in which PKCẟ^+^ neurons are fluorescently labeled with tdTomato, we detail the distinct morphological features of the PACAP-immunoreactive axonal terminals in the CeC, particularly enriched in its postero-ventro-lateral region (Fig. 1 C). These terminals display a perisomatic “calyx-like” arrangement, as seen in high-magnification confocal images (Fig. 1, C1), and juxtapose with GluD1-immunolabeled domains of PKCẟ expressing neurons (Fig. 1, D), supporting a direct axo-somatic PACAP containing input onto PKCẟ^+^ /GluD1^+^ neurons.

### Electrophysiological analysis of PACAP-driven responses

In order to determine the postsynaptic response of PKCδ-expressing neurons in the CeC to PACAP, we performed *ex vivo* whole-cell patch-clamp recordings in horizontal brain slices from Prkcd-Cre::Ai9 mice, in which PKCẟ^+^ neurons are labeled with tdTomato. Neuronal excitability was quantified using a repeated current-step protocol (1s steps from -40 to +240 pA in 20pA increments, 7 s intervals) applied during three 20 min conditions: baseline aCSF (artificial cerebrospinal fluid), PACAP (100 nM), and washout. The protocol ran continuously, and five runs per condition were averaged; for PACAP and wash, only runs obtained after the first 10 min of each condition were analyzed. Current-frequency (I-F) curves were generated for each neuron, and a two-way repeated-measures ANOVA (condition × current step) was used to determine PACAP effects on excitability. Neurons showing a significant increase (p<0.05) in firing during PACAP were classified as type I (PACAP-responsive), whereas those without significant changes were classified as type II (non-responsive). In total, 30 PKCẟ^+^ capsular CeA neurons were recorded and assigned to one of these two phenotypes. Type I neurons (symbolized with red crosses in Fig. 2, panel A, n=21, located in the ventro-lateral segment of CeC) show a robust and reversible increase in firing rate during PACAP application (Fig. 2, panel B), along with a steeper current–frequency (I-F) relationship (Fig. 2, C), while type 2 neurons (symbolized with black dots in panel A, n=9) exhibit no significant change in excitability (Fig. 2, E). For PACAP-responsive neurons (type I), a two-way ANOVA revealed significant main effects of treatment, F(2,780) = 25.64, p < 0.001, and current injection steps, F(12,780) = 298, p < 0.001, with no significant interaction between the factors. *Post hoc* multiple comparisons confirmed a significant increase in excitability during PACAP application relative to baseline (p < 0.001), with firing rates returning to baseline levels during the washout. Furthermore, *post hoc* immunohistochemistry demonstrated that these PACAP-responsive neurons expressed GluD1 at the plasma membrane (Fig. 2D, left panel), and their morphology was confirmed by biocytin filling (Fig. 2D, right panel). These characteristics are consistent with the PACAP-containing axons perisomatically innervating a subpopulation of PKCẟ^+^ neurons described in Figure 1. Interestingly, long projecting axons of the type I neurons were observed (Fig. 2D, right panel, main axon was indicated by arrows). The type II neurons showed no significant change in firing frequency during PACAP application (Fig. 2E-F). The two-way ANOVA for Type 2 neurons revealed a significant effect of current injection steps, F(12,312) = 84.1, p < 0.001, but no significant effect of treatment, nor any significant interaction between the factors. Additionally, these PACAP-insensitive neurons lacked membrane-localized GluD1 despite being tdTomato-expressing (Fig. 2F, left panel), and their morphology was confirmed via biocytin labeling (Fig. 2F, right panel).

### Molecular signature of calyceal terminals engulfing the PKCδ-ir neurons in CeC

High-resolution confocal imaging revealed that these ring-like axon terminals co-expressed the vesicular glutamate transporter 1 (VGluT1) and/or 2 (VGluT2) (Fig. 3, panel D, A11, 12, A17, B6 y B7 and Fig. 5 panel H, at electron microscopy level), analogously to the calyx of Held in the medial nucleus of trapezoid body (MNTB), and the vesicular acetylcholine transporter VAChT (Fig. 3, panels B). Furthermore, the presynaptic PACAP^+^ rings around the somata of PKCδ-expressing neurons contained the neuropeptides calcitonin-gene related peptide (CGRP) and neurotensin (NTs), as well as the calcium binding protein calretinin (Fig. 3, panels C). These findings indicate a multimodal transmitter phenotype for these axon terminals that include glutamatergic, cholinergic, and peptidergic components. To prove that their spatial organization is similar to that of a calyx of Held (calyx-like), we performed confocal z-stacks of PKCδ-expressing neurons segmentation in the CeC. The PACAP^+^ fibers formed large cup-like, enveloping axo-somatic terminals, confirming their calyx-like macro-architecture (Fig. 3, D).

### Ultrastructure of the extended-amygdala calyceal synapse (EM level)

Serial FIB-SEM tomography (20 × 20 × 20 nm voxels; Fig. 4A and SI Figure 3) of PACAP immunoreacted CeC sample revealed a single, giant presynaptic terminal that engulfs the somatic surface of its postsynaptic neuron. Three-dimensional reconstructions show a cup-like shell perforated by slender filopodial protrusions that track the contour of the target soma (Fig. 4, panel A2, A3 and A4, and SI Figure 3 B). Inside the calyceal lumen, the postsynaptic membrane is smooth and devoid of spines, reminiscent of the principal cell in the classical calyx of Held (Fig. 4, A1, and SI Figure 3, also see the supplemental information videos 1, 2, 3).

**Figure 4.**
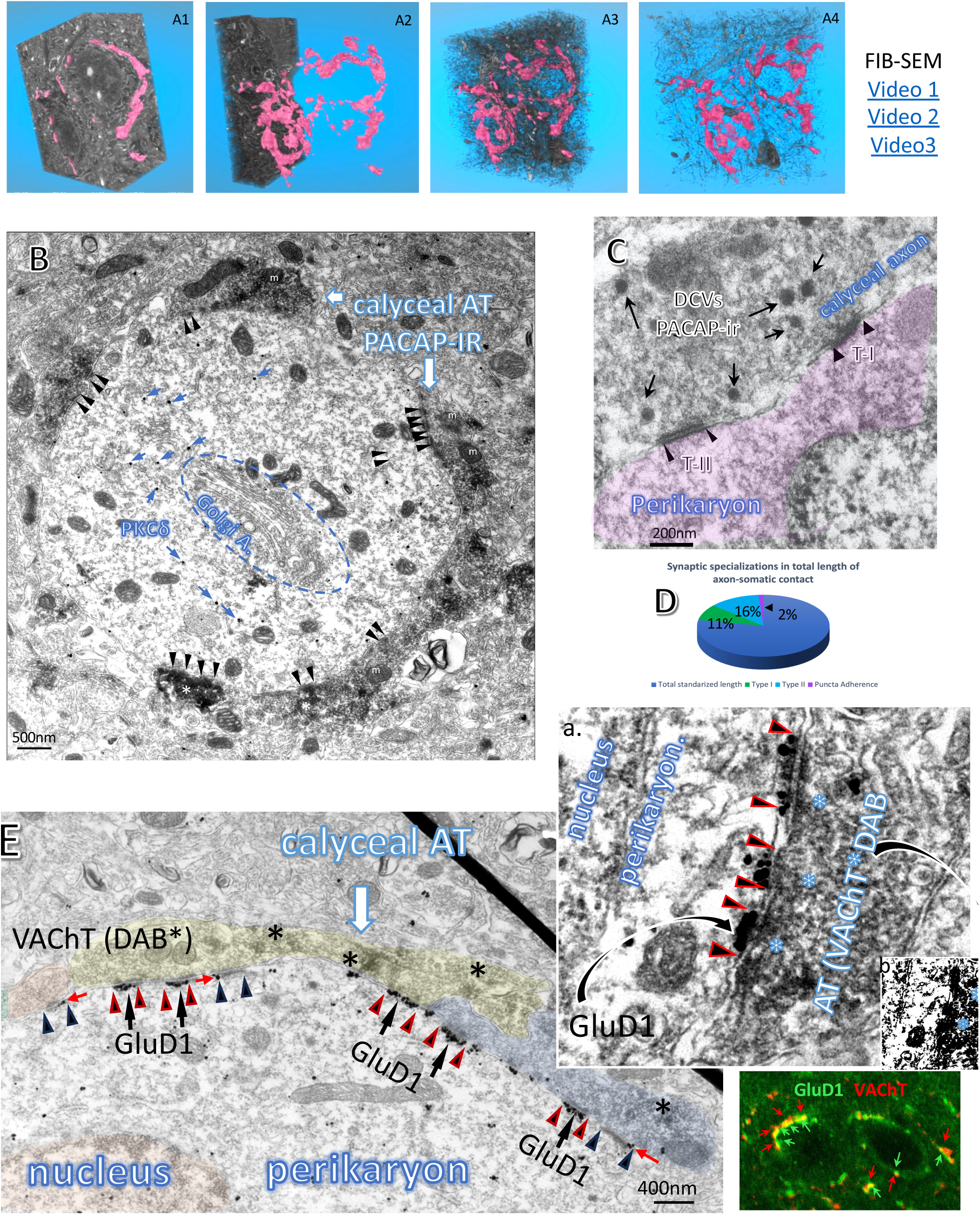
Electron microscopy showing distinct features of calyceal synapse in the extended amygdala. **(As)** A calyx of Held-like, ring-shaped synaptic structure in the central amygdala (CeC-calyceal synapse) revealed by immuno-electron microscopy against PACAP FIB-SEM (Focused Ion Beam Scanning Electron Microscopy). **(A1-A4)** 3D reconstruction and segmentation of the same sample highlighting the ring-like morphology of the PACAP^+^ calyceal synapses. Corresponding video files showing the full reconstruction are available in the Supplemental Information videos 1, 2 and 3. **(B)** a double-immuno-electron microscopy photomicrograph showing a PACAP-ir calyceal axon terminal forming numerous synaptic specifications (arrowheads) onto a PKCẟ labeled (gold particles) soma (a Golgi apparatus was indicated). **(C)** A PACAP-expressing calyceal axon forms both asymmetric (type I, thick arrowheads) and symmetric (type II, narrow arrowheads) synaptic contacts. Presynaptic vesicles and both pre- and postsynaptic densities (PSDs) are visible. Arrows indicate PACAP^+^ dense core vesicles. **(D)** A pie chart quantifies the proportion of each synapse type based on contact length measurements from randomly selected TEM micrographs (n = 30) in which synaptic specializations could be identified. **(E)** Another example of a VAChT^+^ (* DAB) calyceal axon forming Type II synapses with GluD1-gold labeling at the post-synaptic density (PSDs). Note that the GluD1 gold particles are selectively located at the PSDs of type II synaptic specifications (red-black arrowheads). In contrast, the PSDs of 3 adjacent type I (asymmetric) synaptic specifications (black arrowheads) lack the GluD1 labeling. Top inserts: “a” shows higher magnification of this peculiar type of cholinergic synapse with DAB immunolabeling against VAChT at the active zone (indicated by asterisks) and the post-synaptic densities immunopositive to GluD1 (flanked by arrowheads). Insert “b”: over-contrasted “a” to show the DAB deposition in the axon terminal. Lower insert: confocal image showing close spatial correspondence between VAChT (red) and GluD1 (green) signals. GluD1 expression is localized between the red VAChT-positive axon and the soma, suggesting a tight anatomo-functional association between cholinergic input and GluD1 expression.

At higher magnification (Fig. 4, B,C), multiple synaptic specializations (SS) can be observed within the calyceal structure (indicated with arrowheads). The active zones of those SS are packed with membrane bound clear, round vesicles and scattered large dense-core vesicles (indicated by white asterisks) that immunoreacted for PACAP. Mitochondria (“m” in panel B) are abundant within the calyceal structure. We revealed that PACAP-ir calyceal terminal formed both asymmetric (type I) and symmetric (type II) synaptic contacts (Fig. 4C), a higher prevalence of type II configurations (Fig. 4D). This type II SS showed strong deposition of electron dense materials in the presynaptic active zone coincided with vesicular Ach transport (VAChT) immunolabeling (Fig. 4E, “*” indicate DAB deposition linked with VAChT-ir. The inset “a” of panel E shows higher magnification of this peculiar type of cholinergic synapse with DAB immunolabeling against VAChT at the active zone (indicated by asterisks) and the post-synaptic densities immunopositive to GluD1). Inset “b” of panel E was overtly contrasted to show the DAB deposition at the axon terminal. The immunofluorescent inset shows double immunofluorescence that GluD1 (green) labeled plasma membrane apposed with VAChT-ir patches in the outline of the cell (red). Hence, these are symmetrical SS of cholinergic type II synapses (20, 21). Noteworthy, we often detected strong GluD1 immunogold labeling of the postsynaptic density (PSD) of the type II SS. Figure 4 panel E show an example of this calyceal synapse with mixed SS. The black arrowheads flank the type I SS, while the black arrowheads-outlined with red line flank the type II SS. Strikingly, the GluD1 immunolabeling (gold particles, indicated with black arrows) are selectively deposited in the PSD of the type II SS and only very few of them are deposited at the edge of the type I SS (red arrows) suggesting a specific molecular identity for this synaptic pairing. In SI Figure 4, we show more examples (panels C, D, E, F and G) reinforcing the idea that GluD1 expression is closely associated with cholinergic inputs. The Glu1D has been interpreted as an adhesion molecule (22, 23) in recent literature. Hence an interesting question arises here that if this is the case, why is this selective distribution? These findings support the notion that CeC-calyceal synapses represent a specialized subpopulation of central amygdalar connections, distinguished by their molecular profiles and potential neurotransmitter signaling mechanisms.

### Brain-stem origin of the calyceal input

Finally, to determine the origin of the calyx-like PACAP^+^ terminals, we performed tract-tracing experiments. Stereotaxic deposits of the retrograde tracer fluoro-gold (FG) confined to the CeC (Fig. 5A) labeled a discrete neuronal cluster of large cells in the ventrolateral parabrachial complex (PBc), centered in the Kölliker-Fuse (KF) nucleus (Fig. 5B). Most retrogradely labeled neurons in this region co-expressed *Slc17a7* (VGluT1), *Slc17a6* (VGluT2), and *Adcyap1* (PACAP) mRNAs (Fig. 5), as detected by multiplex RNAscope method, indicating a glutamatergic/PACAP phenotype that matches the neurochemistry of the calyceal terminals identified in the EA. In SI Figure 5, we present six combinatory dual *in situ* hybridization (DISH) experiment that revealed the large cells in the Kölliker-Fuse area co-expressed mRNAs of *Adcyap1* (PACAP) with *Slc17a7* (VGLUT1) (A), with *Slc17a6* (VGLUT2) (SI Figure 4, B), with *Chat* (CHAT) (SI Figure 4, C), with *Calca* (CGRP) (SI Figure 4, D), as well as *Slc17a7* (VGluT1) co-expression with *Slc17a6* (VGluT2) (SI Figure 4, E) and with *Chat* (CHAT) (SI Figure 5, F).

**Figure 5.**
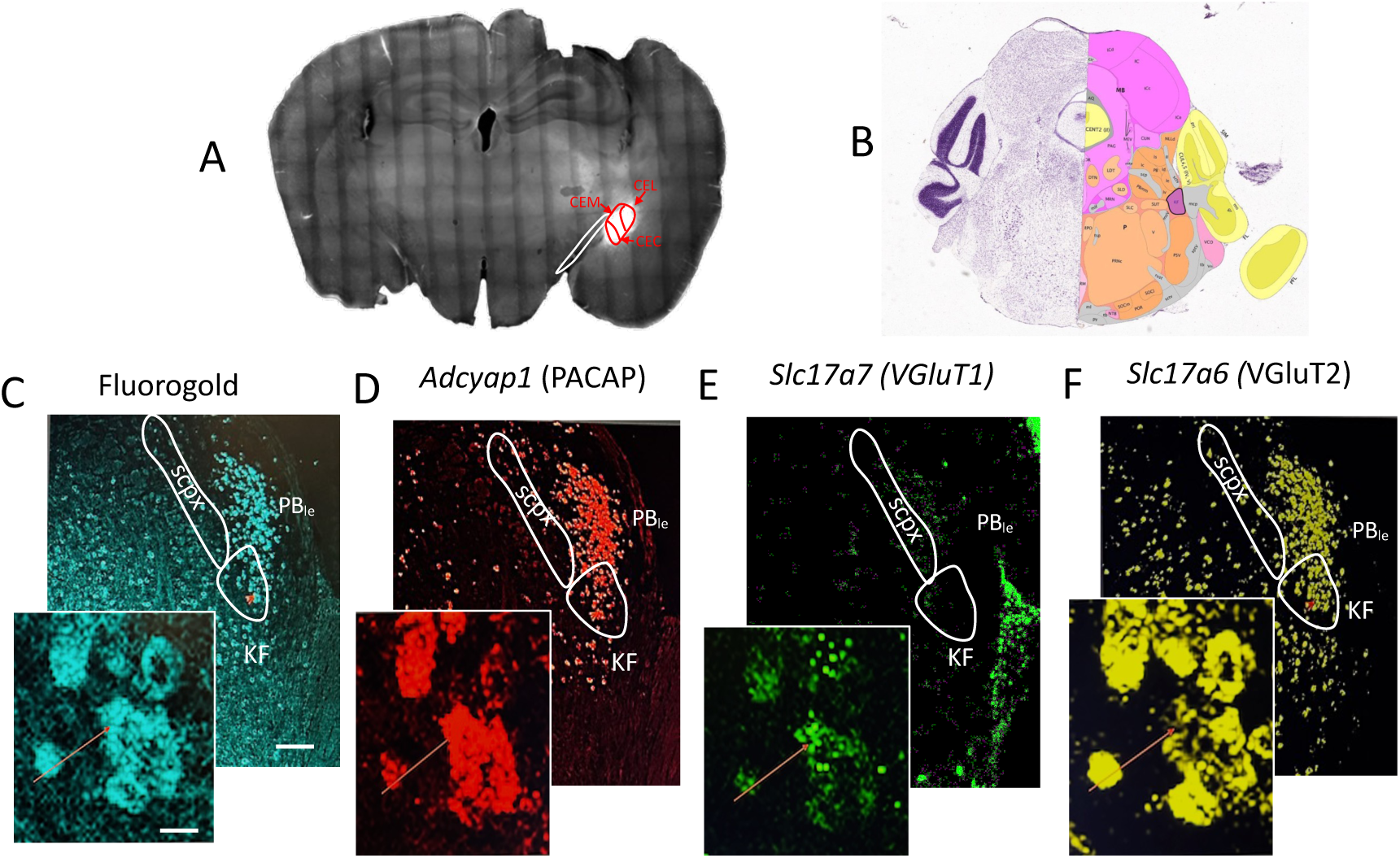
Retrograde labeling with fluorogold injected within mouse central amygdala reveals the pontine Kölliker–Fuse (K-F) subnucleus hosts the input neuronal population with molecular signature corresponding the calyceal synapse. **(A)** Fluorogold was precisely injected into central amygdala (red outlines; optic tract indicated with white outline) two weeks before the multiplex in situ hybridization using RNAscope multiplex method. **(B)** Mouse pons coronal section showing the location of the KF Koelliker-Fuse subnucleus (dark pink shadowed area) within the parabrachial complex. **(C)** Fluorogold (FG) found in the parabrachial complex, especially in the latero-external subdivision (PBle and Kölliker-Fuse subnucleus (KF) overlapping with (D) *Adcyap1* (PACAP), (E) *Slc17a7* (VGluT1) and (F) *Slc17a6* (VGluT2). The insets with confocal microscope software general arrows indicating the same cells. Scale bars: low magnifications: 100µm, high magnification: 20µm.

To further prove that PACAP^+^ neurons in the PBc give rise to calyx-like terminals, we injected a low volume of a Cre-dependent AAV5-EF1α-DIO-hChR2(H134R)-eYFP into the KF of PACAP-Cre mice (Fig. 6A). Despite sparse transduction of only a handful of PACAP^+^ somata per case (Fig. 6B), eYFP+ axons produced stereotyped ring-like endings in both CeC and BSTov (Fig. 6 C and D), recapitulating the perisomatic geometry seen with immunohistochemistry (respective insert showing high magnification of calyceal innervations). In a parallel experiment, we employed freeze-fracture replica immunolabeling (FRIL) (24) to visualize the eYFP+ inputs in the CeC. Large membrane patches of axonal calyceal innervation displaying a high density of immunogold particles for ChR2-eYFP were observed over the somatic membrane of postsynaptic neurons (Fig. 6 G), consistent with enveloping calyx-like structures. Accumulations of immunogold particles detecting the extrasynaptic domain of AMPA receptors were also observed on the somatic exoplasmic-face (E-face) opposite of the eYFP-labelled protoplasmic-face (P-face) membrane (Fig. 6 panels E and arrowheads in G), suggesting the presence of small glutamatergic synapses, despite no clear clusters of intramembrane particles, typically defining type I postsynaptic specializations, could be identified. Typical axon terminals (at) forming synapses with dendritic spines (sp) was not found to display labelling for ChR2-eYFP (Fig. 6 F), further supporting the evidence that transduced KF neurons specifically form calyx-like terminals.

**Figure 6.**
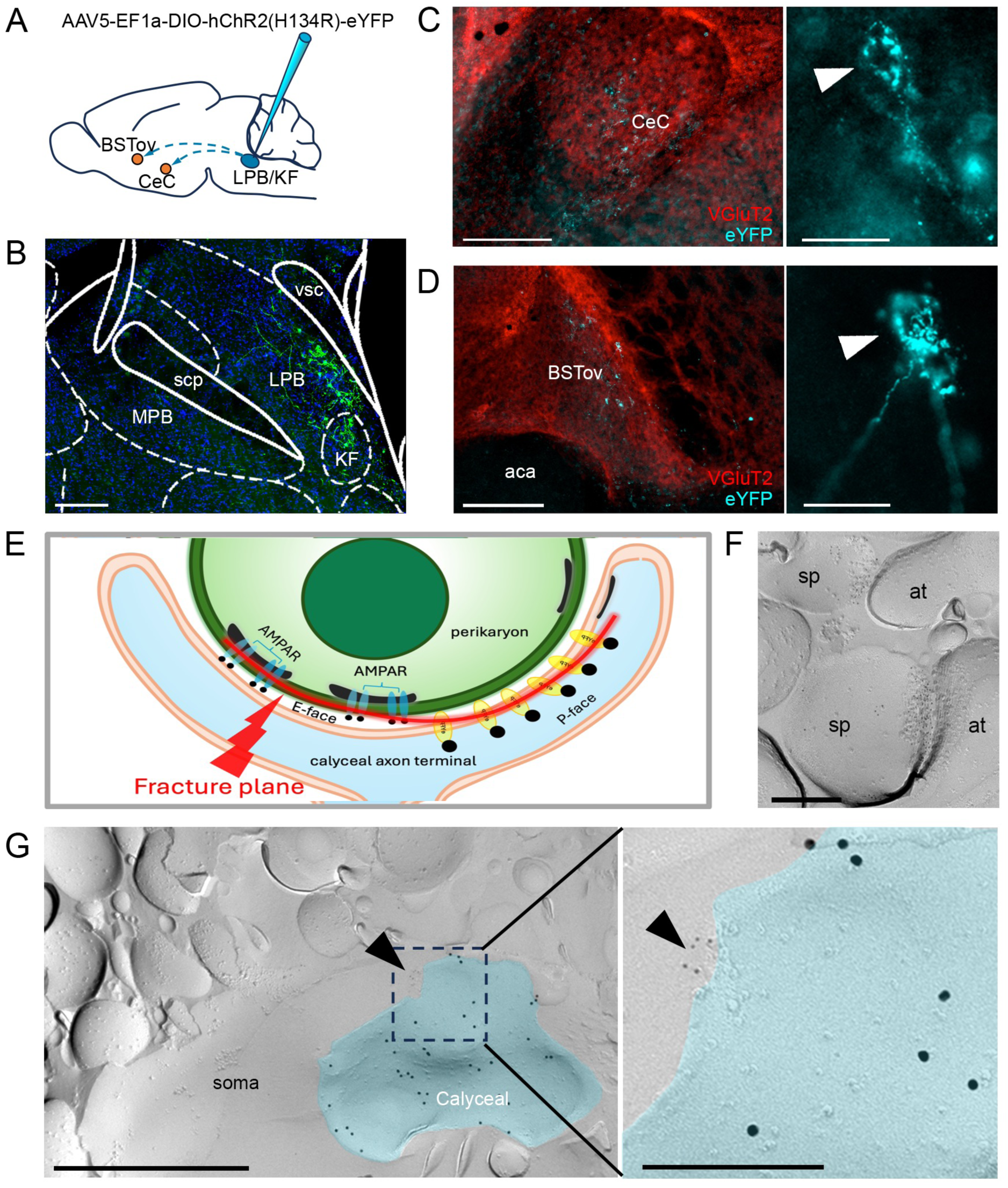
Axons originated from the KF-LPB give rise to calyx-like terminals which form putative synaptic contacts with the soma of CeC neurons containing AMPA receptors as revealed by Freeze-Fracture Replica Immunolabeling (FRIL). (**A**) Schematic drawing of the AAV5-EF1a-DIO-hChR2(H134R)-eYFP injection site. (**B**) Micrograph taken from a representative coronal section of the pons showing ChR2-eYFP-transduced neurons of PACAP-cre mice (n=13), confirming that the microinjections were correctly placed into the KF-LPB. DAPI staining (blue) was used to reveal the macrostructure of the brain slice. Scale bar: 200 µm. (**C)** Micrographs of the KF-LPB efferent projections (turquoise) in the capsular subdivision of the central amygdala (CeC) and (**D**) in the oval nucleus of the bed nucleus of the stria terminalis (BSTov). The sections were co-stained for VGluT2 (in red). Scale bars: 250 µm. Panels on the right of C and D are higher resolution micrographs showing calyceal innervation from PACAP-neurons of the KF-LPB expressing eYFP, in CEC and BSTov, respectively. Scale bars: 50 µm. (**E**) Schematic drawing summarizing the spatial organization of presynaptic and postsynaptic elements. Upon fracture (red line symbolize the fracture plane), the presynaptic membrane of the calyceal axon terminal is exposed as a P-face containing numerous 15 nm immunogold particles identifying ChR2-eYFP. The complementary E-face of the postsynaptic somatic membrane displays discrete clusters of 5 nm gold particles marking AMPA receptors. (**F**) Axon terminals (at) forming synapses with spines in the CeC did not display immunolabelling for ChR2-eYFP. The postsynaptic specializations, visible as an intramembrane particles (IMP) accumulation on the spine head, show a clear accumulation of AMPA receptors (5 nm gold particles). Scale bar: 250 nm. (**G**) Electron-micrograph of a CeC replica. Scale bar: 1 µm. Labelling for ChR2-eYFP (large gold particles, 15 nm) identifies the protoplasmic face (P-face) of the calyx-like terminal originating from the KF-LPB; 5 nm gold particles (arrowheads) reveal AMPA receptors at the exoplasmic membrane (E-face) of the perikaryon of CeC neurons. A pseudocolor is used to simplify the identification of the P-face of the calyx-like membrane, identified by the presence of numerous 15 nm gold immuno-particles against ChR2-YFP. The inset shows a higher magnification of the squared region. No obvious accumulation IMP could be observed associated with the cluster of 5 nm gold particles. Scale bar: 100 nm. Abbreviations: aca, anterior commissure, anterior part; BSTov, bed nucleus of the stria terminalis, oval subnucleus; CeA, central nucleus of the amygdala; KF, Kölliker-Fuse nucleus, LPB, lateral parabrachial nucleus; MPB, medial parabrachial nucleus; scp, superior cerebellar peduncle; VGluT2, vesicular glutamate transporter 2; vsc, ventral spinocerebellar tract.

Together, these results identify PACAP/VGluT1/VGluT2 KF neurons as a sparse presynaptic population capable of generating calyx-like terminals in the EA. These anatomical data align with our rat study using hydralazine-evoked hypotension, where a single juxtacellularly labeled KF neuron sent a rostral axon through the superior cerebellar peduncle (scp) to dorsal tegmental bundle to the CeC, and continued via the *ansa peduncularis* to the BSTov, forming ring-like (calyceal-like) perisomatic terminals in both targets ((25), in their Fig. 5). Thus, convergent retrograde, molecular, and single-cell tract data point to the KF as a principal source of the EA calyces.

## Discussion

Giant axo-somatic excitatory synapses are exceedingly rare in the mammalian brain and, until now, have been confined to the auditory brainstem, where the calyx of Held ensures *high-fidelity* transmission required for precise sound localization (26, 27). Here we identify a multimodal calyceal synapse in the extended amygdala (EA) - specifically in the capsular central amygdala (CeC) and oval bed nucleus of the stria terminalis (BSTov) - revealed at the electron microscopic level through serial FIB-SEM reconstruction and correlated transmission immuno-electron microscopy. This forebrain calyx-like terminal, originating from PACAP-expressing neurons in the pontine Kölliker–Fuse (KF) nucleus, represents a previously unrecognized synaptic organization for rapid viscerosensory drive into limbic-autonomic circuits. Its giant, enveloping structure, transmitter diversity, and precise postsynaptic targeting distinguish it from all known forebrain or limbic synapses. Complementing the structural analysis here, a companion study shows that this KF nucleus PACAP/VGluT1 pathway is selectively engaged immediately after the acute hypotension challenge, preceding a delayed magnocellular SON/PVN activation (25).

### Multimodal transmitter convergence in a single giant terminal

The KF-derived calyces co-express VGluT1/2 (glutamate), VAChT (acetylcholine), PACAP, CGRP, neurotensin, and calretinin, forming mixed type I and II synaptic specializations around their target somata. This represents an unprecedented level of transmitter convergence within one presynaptic element. In other brain regions, dual co-transmission (e.g., glutamate with ACh in habenular or basal forebrain neurons) supports temporal complementarity of fast ionotropic and slower metabotropic signaling, where distinct transmitter classes are packaged in separate vesicle types and released depending on neuronal activity and intracellular calcium dynamics (28, 29) (30, 31). The combination of multiple peptides in a single axo-somatic terminal implies a broader functional bandwidth. However, it remains to be tested.

### GluD1 likely plays a roles beyond adhesion

In the postsynaptic side, the calyces selectively target PKCẟ⁺ neurons that co-express GluD1, an atypical glutamate receptor homologue originally considered a synaptic adhesion molecule (32, 33). However, recent studies have redefined GluD1 as a metabotropic signal transduction device that couples to intracellular second messenger cascades and regulates synaptic organization and long-term potentiation (22, 23). In our material, GluD1 immunogold particles were concentrated specifically at the symmetric (type II) synaptic contacts apposed to VAChT⁺ presynaptic domains, but were largely absent from the asymmetric (type I), glutamatergic synapses within the same calyx (Fig. 4E). This highly selective distribution suggests that GluD1 participates in cholinergic and peptidergic synapse specialization, potentially coordinating postsynaptic responses to combined ACh/PACAP signaling.

### A specialized Kölliker–Fuse subpopulation as the origin of forebrain calyces

Using retrograde and anterograde tracing we pinpointed the origin of these multimodal calyces to a discrete neuronal cohort in the Kölliker-Fuse subnucleus of the parabrachial complex, distinguished by co-expression of VGluT1, VGluT2, and PACAP mRNAs. The KF is a functionally heterogeneous structure involved in respiration, arousal, and cardiovascular regulation (34–36), but this particular subpopulation of PACAP/VGluT1/2⁺ neurons is exceptionally well positioned to broadcast visceral state information to higher centers. The coexistence of glutamatergic and peptidergic machinery enables both rapid, time-locked excitation and slower, state-dependent modulation. Our combined viral tracing and FIB-SEM evidence demonstrates that these KF neurons generate the giant axo-somatic calyces in both CeC and BSTov, suggesting a unified pathway for high-fidelity transmission from the pons to limbic forebrain. The discovery of this KF lineage implies that calyceal synaptic precision is not restricted to the auditory pathway but can be redeployed in viscerosensory–emotional circuits. By broadcasting visceral state information through axo-somatic calyces, KF neurons may ensure that extended-amygdala ensembles receive temporally precise yet modulatable input, enabling rapid synchronization of defensive or homeostatic responses across CeC and BSTov.

### Functional implications and circuit integration

Although our study applied mainly anatomical methods, the structural organization of this ponto-limbic pathway offers an intuitive functional logic. The KF to EA calyx likely provides a high-gain, short-latency input capable of driving PKCẟ⁺ neurons synchronously in response to visceral perturbations and aligns with the observed two-tier temporal sequence during hypotension - early KF to EA activation for sympathetic rebound and later SON/PVN recruitment for neurohormone mediated stabilization (37).

Within the extended amygdala, PKCẟ⁺/ GABAergic neurons in CeL/CeC inhibit GABAergic output neurons in medial CeA (CeM), establishing a local disinhibition motif that gates downstream autonomic pathways (38). Through anterograde tracer and electron microscopy (EM), CeM neurons have been shown to monosynaptically innervate the rostral ventrolateral medulla sympathetic center (RVLM), including appositions onto phenylethanolamine-N-methyltransferase-positive adrenalin-synthetising neurons that were Fos-positive after hypotension (39). Moreover, CeA and BNST GABAergic projections modulate PVN activity indirectly via the peri-PVN GABAergic zone, which provides tonic inhibition to hypothalamic paraventricular nucleus (PVN) via a peri-PVN GABAergic shell hence is predicted to disinhibit PVN (40). In turn, PKCẟ⁺ neurons in CeC and BSTov, known to be GABAergic and to project to preoptic and hypothalamic inhibitory relays, may exert disinhibitory control over presympathetic neurons in the paraventricular nucleus (PVN) (18, 41).

### Concluding remarks

By uncovering a multimodal calyx-like synapse in the forebrain, this study broadens the known repertoire of mammalian synaptic architectures and demonstrates that calyceal precision is not exclusive to sensory processing. The structural integration of glutamatergic, cholinergic, and peptidergic signaling within a single giant synapse suggests a mechanism by which the forebrain can couple rapid neural responses to modulatory context, uniting visceral feedback with emotional and homeostatic regulation. Future work should examine whether similar multimodal calyces exist in other limbic or hypothalamic regions and explore how GluD1-mediated signaling contributes to their function and plasticity. Together, these findings redefine the extended amygdala as a high-fidelity viscerosensory hub and establish a framework for understanding how the brainstem orchestrates emotional and autonomic integration at the cellular level.

## Materials and Methods

All procedures were conducted in accordance with institutional and national guidelines for the care and use of laboratory animals. Experiments in rats were approved by the Ethics Committee of the School of Medicine, UNAM (protocols CIEFM-079-2020 and CIEFM-083-2025). Experiments in mice were approved by the NIMH Animal Care and Use Committee (ACUC protocol LCMR-08).

### Rats

Adult (>60 days) male Wistar rats were obtained from the UNAM School of Medicine vivarium and maintained under controlled temperature and humidity on a 12-h light/dark cycle (lights on 07:00–19:00) with *ad libitum* access to food and water.

### Mice

Adult (>P60) male and female C57BL/6J wild-type mice, PKCδ-Ai9 reporter mice, and Adcyap1-2A-Cre mice (PACAP-Cre; JAX #030155) were used. Animals were housed under standard conditions.

### Animal allocation

A summary of animals used is provided in SI Table 1.

### Immunohistochemistry for light microscopy

Rodents were perfused transcardially with saline followed by 4% paraformaldehyde. Vibratome sections (70 µm) were incubated with primary antibodies against PACAP, VGluT1/2, VAChT, CGRP, neurotensin, and calretinin, followed by fluorophore-conjugated secondaries. Images were acquired using a Leica Stellaris confocal microscope. Full method description and antibody list (sources/dilutions) are provided in SI Methods and SI Table 2.

### RNAscope multiplex *in situ* hybridization

Fresh-frozen 12 µm cryosections were processed using RNAscope multiplex fluorescence (ACDbio) to detect *Adcyap1, Slc17a7, Slc17a6, Adcyap1r1, Grid1, Chrna7, Prkcd,* and *Sst* transcripts. Hybridization, amplification, and imaging followed ACD protocols. Detailed probe information and procedures are described in SI Methods.

### *Ex-vivo* whole-cell electrophysiology

Horizontal brain slices (300 µm) containing the central amygdala were prepared from Prkcd-Cre::Ai9 mice. PKCẟ⁺-positive neurons labeled with tdTomato in the CeC were recorded in current-clamp mode using K-gluconate internal solution. Neurons were exposed to PACAP (100 nM) during repeated current-step protocols (−40 to +240 pA). PACAP-responsive neurons were defined by a significant increase in firing rate (two-way RM-ANOVA). Full slicing solutions, recording conditions, and statistical details are provided in SI Methods.

### Transmission electron microscopy

Forty-micron vibratome sections were processed with silver-enhanced Nanogold immunolabeling for PACAP, VGluT1/2, VAChT, GluD1, and PKCẟ. Ultrathin sections (60 nm) were imaged on a Tecnai G2 12 TEM. Synaptic specializations were classified as asymmetric (type I) or symmetric (type II), and immunogold labeling was quantified across serial sections. Complete EM protocols appear in SI Methods.

### FIB-SEM tomography

PACAP-immunoreacted blocks were resin-embedded and imaged with a ZEISS Crossbeam 550. Serial images were acquired (20 × 20 × 20 nm voxels) after protective platinum coating and fiducial marking. Image stacks were aligned and reconstructed in Dragonfly (ORS). Acquisition parameters and segmentation workflow are provided in SI Methods.

### Freeze-fracture replica immunolabeling (FRIL)

High-pressure-frozen slices containing CeC were fractured, replicated, and immunogold-labeled for ChR2-eYFP and AMPARs. Replicas were imaged on a Philips CM-120 TEM. A detailed FRIL protocol is included in SI Methods.

### Tract tracing

For anterograde tracing, PACAP-Cre mice received unilateral injections of AAV5-EF1α-DIO-hChR2(H134R)-eYFP into the Kölliker-Fuse area using stereotaxic coordinates (AP −5.2, ML +1.5, DV −3.6 mm). After 4–6 weeks, eYFP-positive fibers were assessed in CeC and BSTov. All surgical details are in SI Methods. Retrograde tracing was performed using Fluoro-Gold deposits restricted to the CeC. Detailed method description can be found in SI Methods.

### Statistics

Electrophysiological analyses used two-way repeated-measures ANOVA with post hoc comparisons (α = 0.05). Additional statistical information is reported in SI Methods.

## Acknowledgements

This work was supported by UNAM-PAPIIT (IG200121 to LZ and RAB), the Mexican Secretaría de Ciencia, Humanidad, Tecnología e Innovación (CIORGANISMOS-2025-92 to LZ; CF-2023-G-243 to LZ, RAB, and RHP; CB-238744 to RAB), and the Intramural Research Program of the NIMH, NIH (ZIAMH002386 to LEE). We acknowledge the PASPA-DGAPA sabbatical fellowships (to LZ and VSH), a Secihti sabbatical fellowship (to LZ), and a Fulbright-García Robles Fellowship (to VSH). We thank Rafael Luján, Luis Miguel García-Segura, Javier deFelipe, Ángel Merchán, Sebastian Schädler, and Claus Burkhart for their guidance in FIB-SEM techniques; to Marisela Morales for introducing us to the NIDA Confocal and Electron Microscopy Core, and to Pedro Segura for technical assistance. LZ wishes to express her gratitude to Peter Somogyi, for hosting her 2019 summer study-stay and for time spent together examining raw material under the microscope, and for the inspiring suggestion that this project might reveal a calyx-of-Held–like synapse. His thoughtful reading of an earlier version of this manuscript is deeply appreciated.

## Supplemental information

**SI Figure 1.**
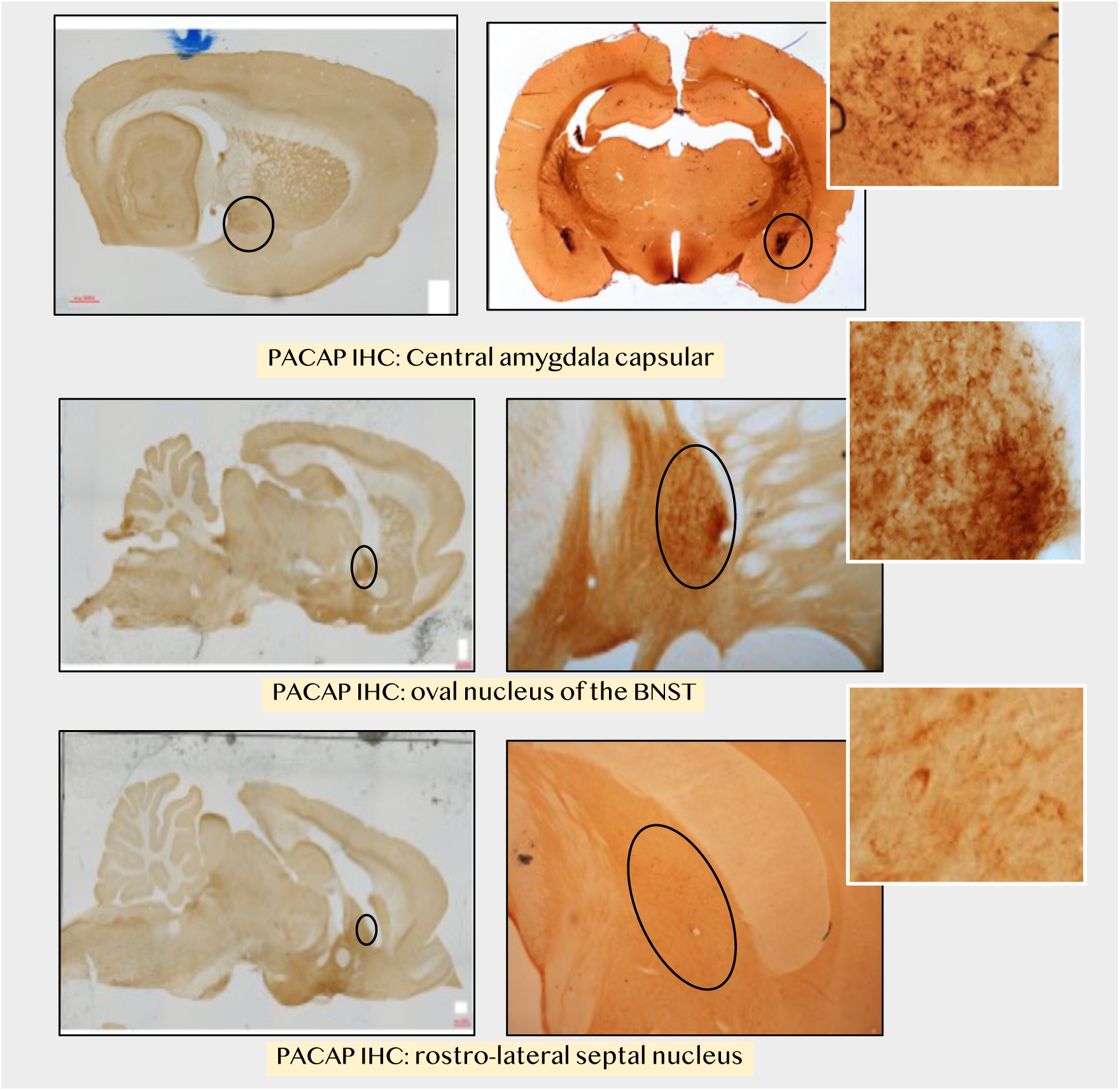
Calyceal axonal endings in male adult rats telencephalic structures observed with PACAP immunohistochemistry. Upper row, central amygdala, capsular subdivision (CeC). Inset is taken from a sagittal section. Middle row: oval nucleus of the bed nucleus of the stria terminalis (BSTov). Bottom row: rostro-lateral septal nucleus.

**SI Figure 2.**
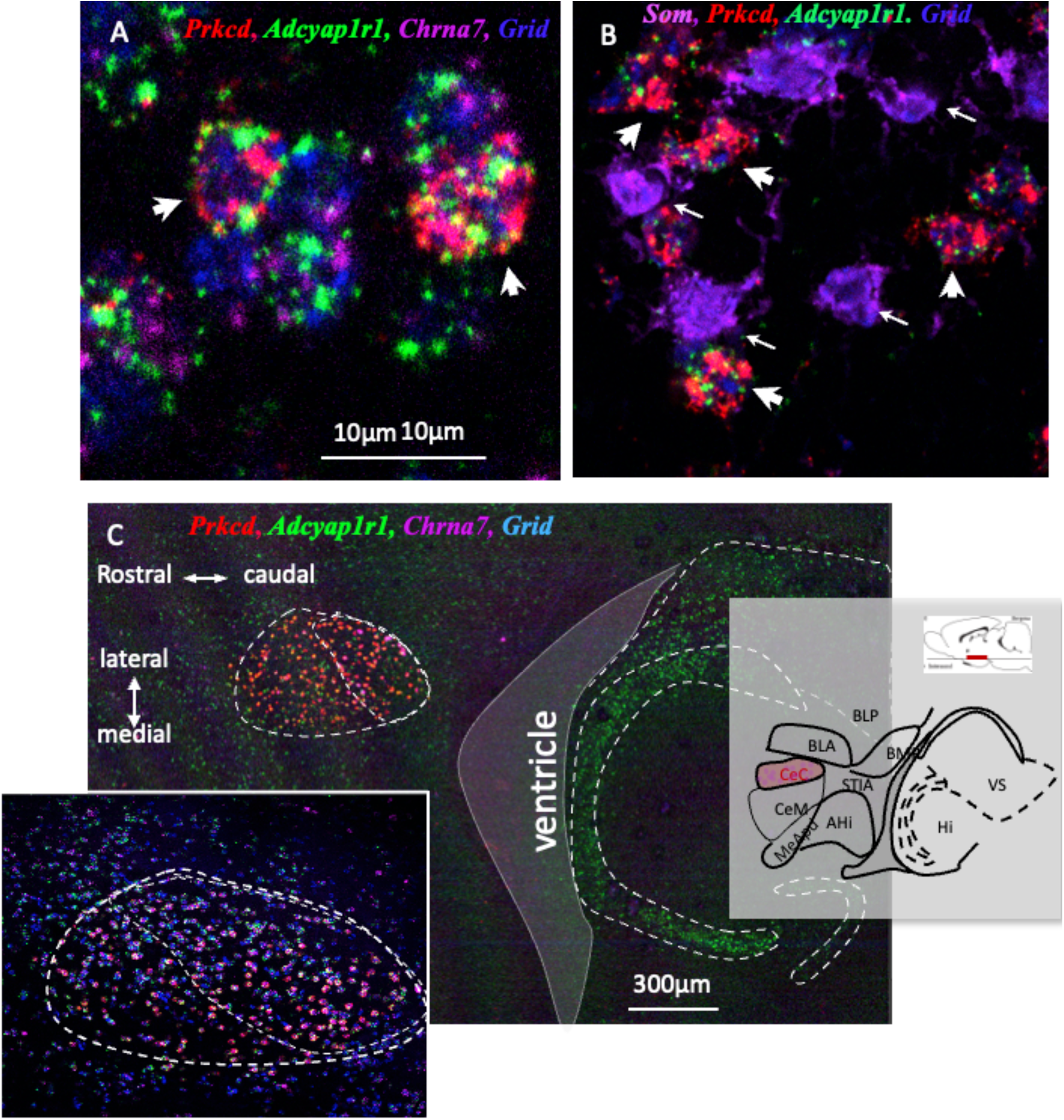
Molecular signature of the calyceal synapse target PKCδ^+^ cells in mouse central amygdala capsular subdivision (CeC). **(A)** *Prkcd* cells co-expressing *Adcyap1r1* (PAC1,) *Grid1* (GluD1), *Chrne* (nAChRa7) (thick arrows); **(B)** *Som* (somatostatin) expressing neurons do not co-express *Adcyapr1r* nor *Prkcd* (thin white arrows). **(C)** Horizontal section showing the selective distribution, in the ventral posterior segment of the CeC of the sub-population of PKCδ^+^ cells *(Prkcd* cells) co-expressing *Adcyap1r1* (PAC1,) *Grid1* (GluD1), *Chrne* (nAChRa7).

**SI Fig. 3.**
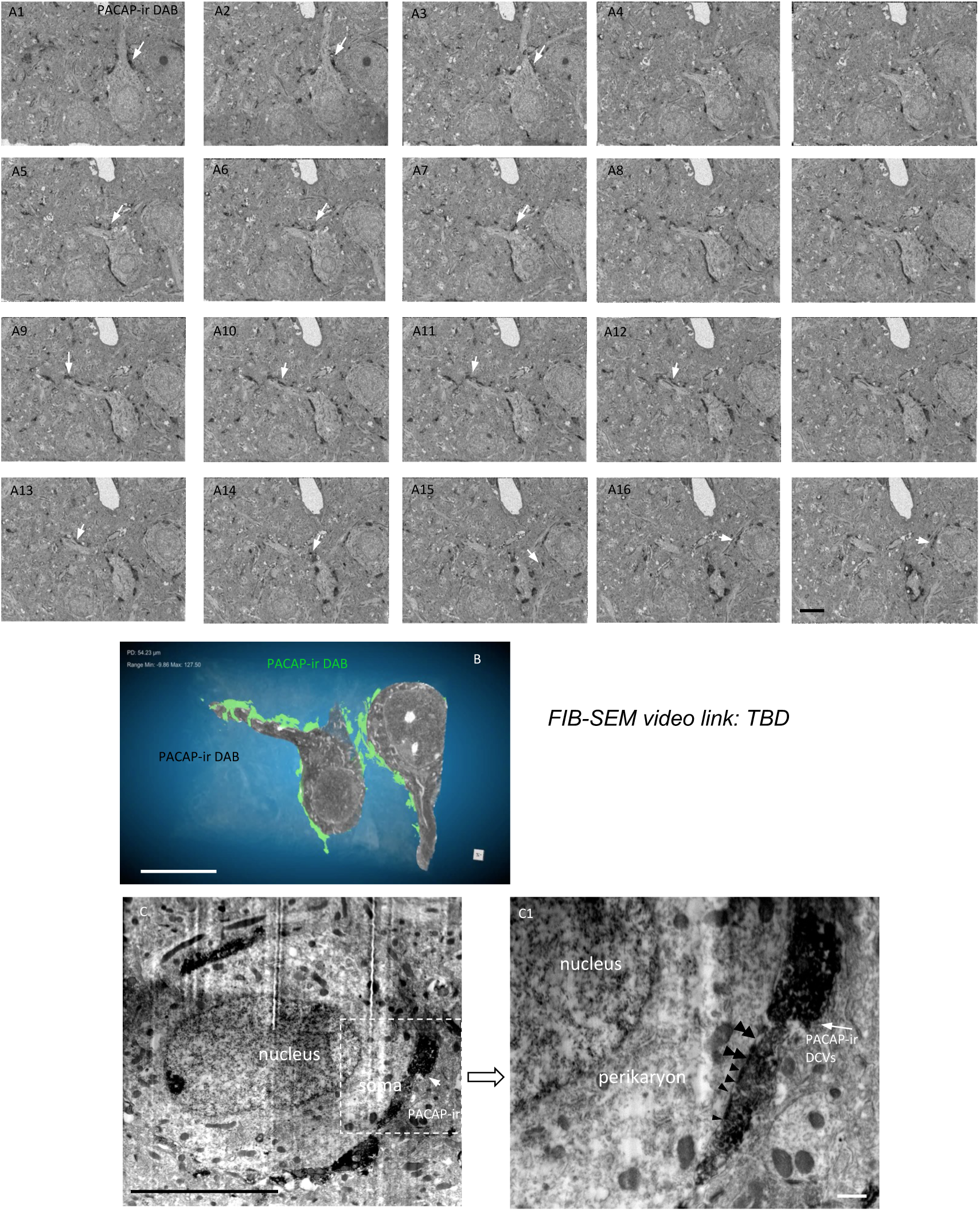
Calyx of Held synapse in extended amygdala under focused ion bean scanning electron microcopy in a PACAP-ir DAB immunoelectron microcopy sample of CeC. (**A1-A17**): Selected frames from a series of 450 focused ion beam scanning electron microscopy (FIB-SEM) images taken from an immunoelectron microscopy sample labeled with rabbit anti-PACAP antibody and visualized using a peroxidase-DAB reaction. These frames illustrate the sequential progression of two PACAP-positive calyceal synapses. **(B)** 3D reconstruction and segmentation of the same sample highlighting the calyx-like morphology of the PACAP+ calyceal synapses Corresponding video files (3) showing the full reconstruction are available in the Supplemental Information. **(C)** a single plane obtained from high resolution FIBSEM scanning showing the PACAP containing axon terminal engulfing the postsynaptic cell body. **(C1)** An amplified segment of the squared region of **C** showing the heterogenous distribution of PACAP-ir large dense core vesicles (DCV, white arrow. Asymmetric and symmetric synaptic specifications limited with double arrowheads and single arrowheads, respectively, are shown. Scale bar: 10µm except C1: 200nm. **Video 1: Computer reconstruction of PACAP immunopositive processes surrounding central amygdala neurons.** **Video 2. Three-dimensional reconstruction of PACAP-immunoreactive processes forming Calyxes with CeA neurons.** **Video 3. Three-dimensional reconstruction of PACAP-immunoreactive calyx-like terminals contacting two neighboring cells.**

**SI Figure 4.**
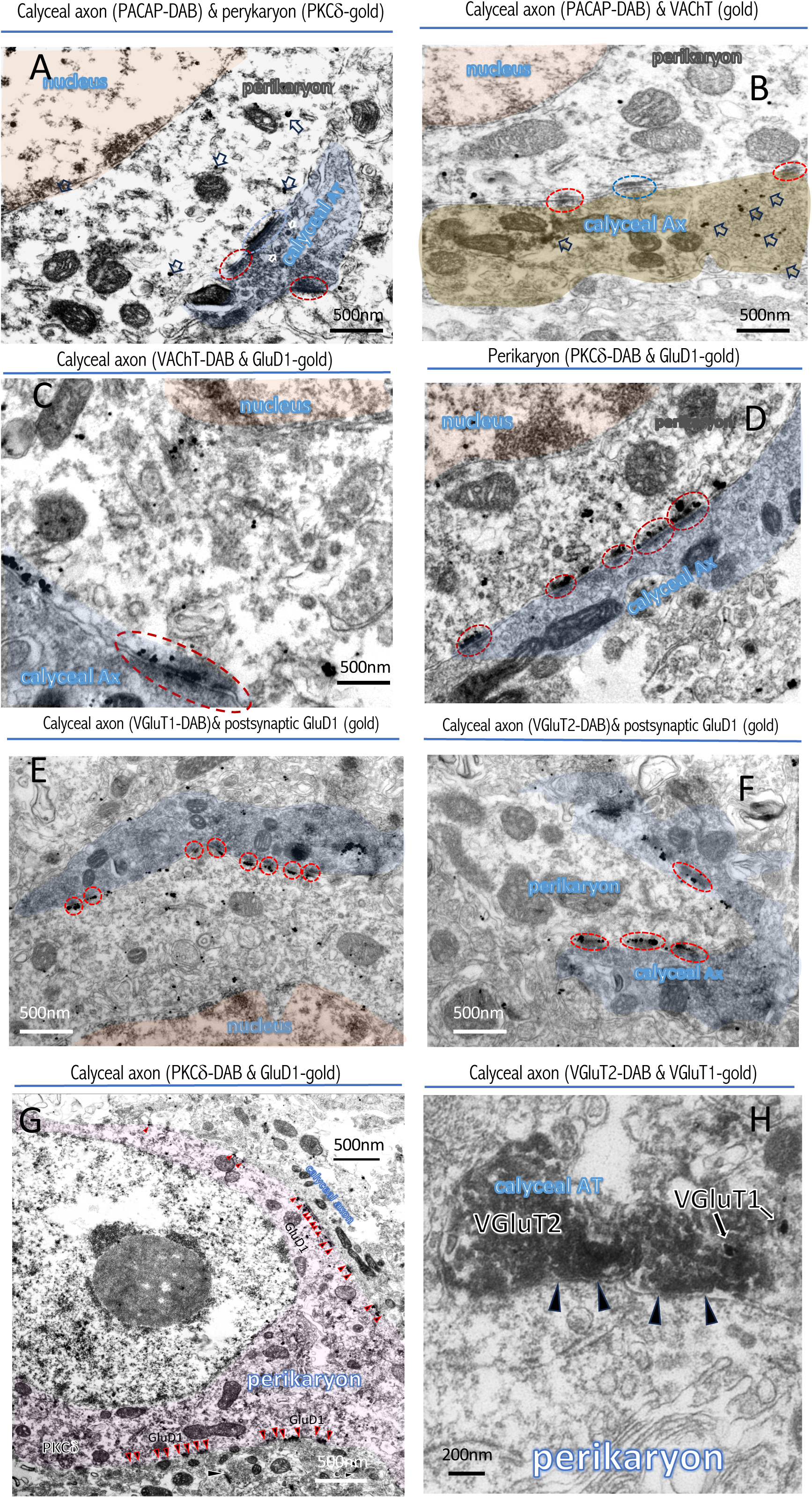
Dual immunoelectron microcopy micrographs showing main ultrastructural features of the calyceal synapse. **(A)** A PACAP-ir calyceal axon (DAB) making type I (blue circle) and type II (red circle) synapses onto PKCδ^+^ labeled soma (gold particle). **(B)** Co-expression of PACAP (DAB) and VAhT (gold particle) of a calyceal axon (DAB) making type I (blue circle) and type II (red circle) synapses onto a soma. **(C)** A VAChT-ir calyceal axon (DAB) making type II (red circle) on to a soma with numerous GluD1-gold particles located at the postsynaptic density (PSD). **(D)** Perikaryon with PKCδ^+^-DAB & GluD1-gold labeling. **(E)** A VGluT1-ir calyceal axon (DAB) with numerous GluD1-gold particles labeled synaptic specifications. **(F)** A VGluT2-ir calyceal axon (DAB) with numerous GluD1-gold particles labeled synaptic specifications. **(G)** A large PKCδ^+^-expressing (DAB-labeled) neuron receives multiple GluD1-positive synaptic contacts (red-black arrowheads) along its soma and proximal dendrite. Two adjacent type I synapses from unlabeled axons are GluD1-negative (white-black arrowheads), underscoring the specificity of GluD1 labeling to type II synapses and suggesting a molecular distinction between glutamatergic and cholinergic inputs. **(G)** VGluT1^+^/2^+^ nature of the calyceal axon terminal. Arrowhead indicate synaptic specifications.

**SI Figure 5:**
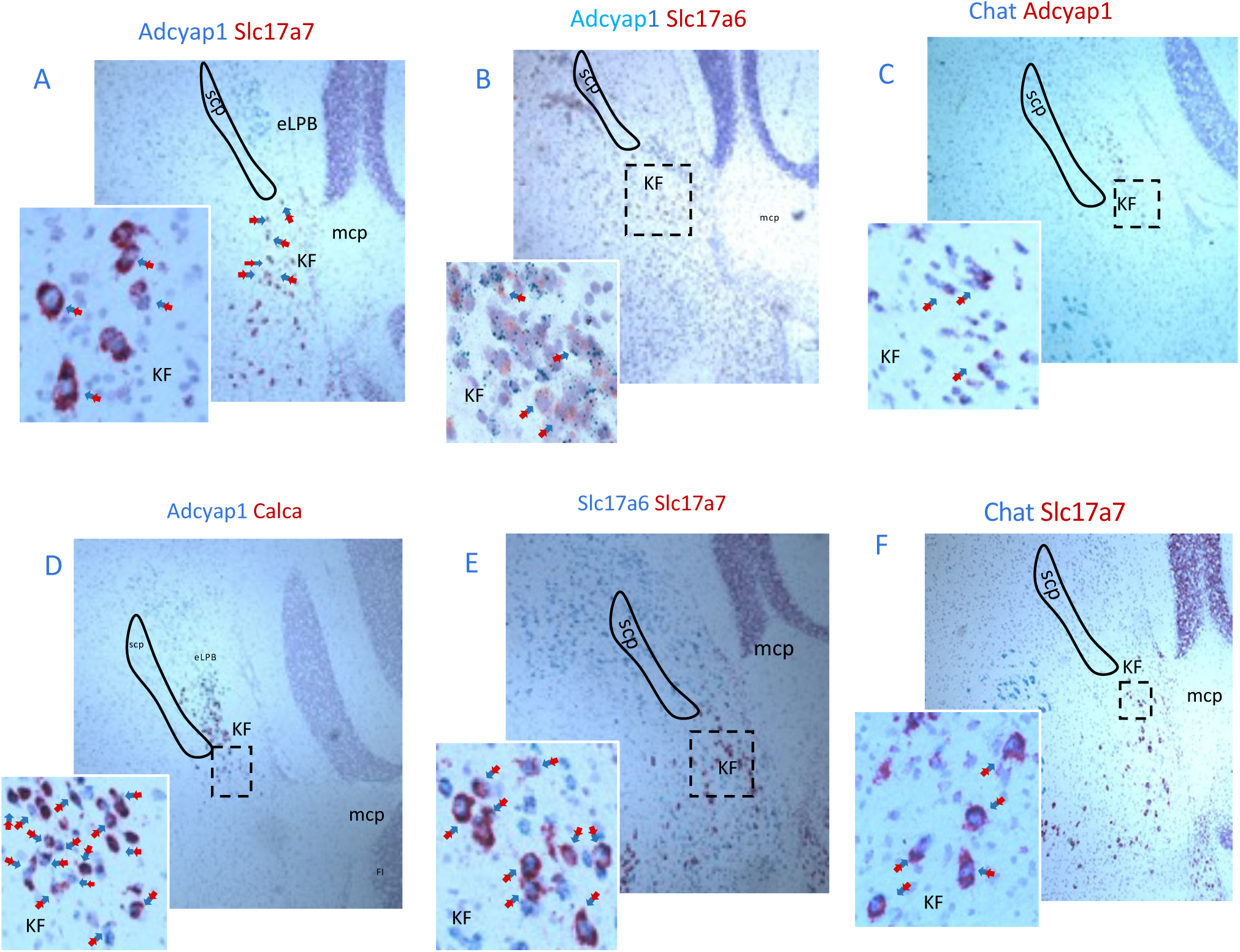
Dual *in situ* hybridization (DISH) revealed the *large cells* in the Kölliker-Fuse area co-expressed mRNAs. **(A)** *Adcyap1* (PACAP) with *Slc17a7* (VGLUT1); **(B)** *Adcyap1* (PACAP) with *Slc17a6* (VGLUT2); **(C)** *Adcyap1* (PACAP) with *Chat* (CHAT); **(D)** *Adcyap1* (PACAP) with *Calca* (CGRP); **(E)** *Slc17a7* (VGluT1) with *Slc17a6* (VGluT2); **(F)** *Chat* (CHAT) with *Slc17a7* (VGluT1).

## Extended Materials and methods

### Animals

#### Rats

Adult (>60 days) male Wistar rats were obtained from the vivarium of the School of Medicine of the National Autonomous University of Mexico. Rats were housed in humidity and temperature-controlled vivarium using a normal light cycle with lights on at 7:00 hours and lights off at 19:00 hours. Animals had *ad-libitum* access to standard laboratory chow and water. All procedures were approved by the Ethics committee of the School of Medicine, of the National Autonomous University of Mexico, license number CIEFM-079-2020 and CIEFM-83-25.

#### Mice

Adult (>postnatal day 60) male and female C57/Bl6J WT, protein kinase C delta - Ai9 (PKCδ-Ai9), and Adcyap1-2A-cre mice (Cre expression restricted to PACAP-expressing neurons of adult were from The Jackson Laboratory (stock # 030155)) were used. Mice were housed in humidity and temperature-controlled vivarium using a normal light cycle with lights on at 7:00 hours and lights off at 19:00 hours. Animals had *ad-libitum* access to standard laboratory chow and water. All procedures were approved by the National Institute of Mental Health Animal Care and Use Committee, NIMH/ACUC approval LCMR-08.

**SI Table 1:** Allocation of animals for each experimental procedure.

| SPECIES / STRAIN | SEX | AGE | EXPERIMENTAL PROCEDURE | N |
| --- | --- | --- | --- | --- |
| WISTAR RATS | Male | adult | Serial Immunohistochemistry for light and transmission/tomographic electron microscopy | 4 |
| PRKCD-CRE :: Ai9 MICE | M/F | adult | Confocal immunohistochemistry and whole-cell patch clamp electrophysiology | 20 |
| C57BL/6J WILD TYPE MICE | M | adult | Immunohistochemistry for light and transmission electron microscopy, fluorogold tracing and, in situ hybridization | 20 |
| PACAP-CRE MICE | M/F | adult | Cre-dependent AAV sparse labelling for anterograde tracing from KF → CeC/BSTov and subsequent freeze-fracture replica immunolabeling | 13 |

### Viral Injection

AAV5-EF1a-DIO-hChR2(H134R)-eYFP (∼4.0 × 10^12^ infectious particles/ml, Lot# 156229) were obtained from UNC virus vector core (Chapel Hill, NC). Surgical procedures and viral injections followed the “NIH-ARAC guidelines for survival rodent surgery.” For tracing or freeze-fracture replica immunolabeling (FRIL), 0.2 µl of AAV5-EF1a-DIO-hChR2(H134R)-eYFP (∼1 × 10^12^ infectious particles/ml, diluted with sterile PBS from stock) were injected slowly (∼0.1 µl/min) into the LPBn, ventral portion (coordinates: AP -5.2 mm, ML 1.5 mm, DV 3.6 mm) of Adcyap1-2A-cre mice, unilaterally. Mice recovered for 4–6 weeks post-injection before undergoing histological verification and processed for FRIL.

### Immunohistochemistry for light microscopy

Rodents were perfused via the ascending aorta with 0.9% NaCl at room temperature, followed by 4% paraformaldehyde. Brain slices (70 µm) were obtained using a vibratome (Leica VT1200). Slices were washed 3 times in PB, blocked in TBST (Trisma Buffer 0.05M, NaCl 0.9%, 0.3% Triton-X) plus 10% Normal Donkey serum for 2 hrs. Slices were then incubated overnight at 4°C with the following primary antibodies: rabbit anti PACAP (1:2000, BMA, T-4473), guinea pig anti VGLUT1 (1:5000, Chemicon, AB-5905), mouse anti VGLUT2 (1:1000, Chemicon, MAB-5504), guinea pig anti VGLUT2 (1:2000, SySy, 135 404), rabbit anti VAChT (produced and provided by Dr. Lee E Eiden, NIMH, NIH (Weihe, Tao-Cheng et al. 1996)), rabbit anti CGRP (1:1000, Phoenix, H-015-09), mouse anti neurotensin (1:1000, Sigma, SAB-4200703), goat anti calretinin (1:2000, Swant, CG1), diluted in TBST plus 1% NDS. The next day slices were thoroughly washed 3 times in TBST and incubated during two hours in the appropriate secondary biotinylated or fluorescent antibodies from Vector Labs at a concentration of 1:500. The processed sections were examined under bright field using a Nikon Eclipse E600 microscope along with its digital camera for photographic documentation and immunofluorescent reactions were observed using a Stellaris Confocal Microscope from Leica Microsystems.

### RNAscope

Brains were rapidly dissected, flash-frozen on dry ice, and stored at −80 °C until sectioning for in situ hybridization (ISH). Coronal brain sections (16 µm thick) were obtained using a cryostat set to −20 °C and mounted directly onto microscope slides. Slides were stored at −80 °C until further processing. RNAscope ISH was performed according to the manufacturer’s instructions (Advanced Cell Diagnostics, ACD) and previously published protocols.

Briefly, slides were fixed in 10% neutral-buffered formalin for 20 min at 4 °C, washed twice with PBS (1 min each), and dehydrated through graded ethanol (50%, 70%, and 100%; 5 min each). Slides were stored in 100% ethanol at −20 °C overnight, then air-dried at room temperature (RT) for 10 min. A hydrophobic barrier was drawn around the tissue sections with a PAP pen and allowed to dry for 10–15 min at RT. Sections were incubated with Protease IV for 20 min at RT, rinsed twice in distilled water (ddH₂O), and hybridized with the appropriate probes for 2 h at 40 °C in a HybEZ™ oven (ACD).

For multiplex experiments, the following probes were used (all from ACD): Mm-Adcyap1 (Cat. No. 405911-C1 and 405911-C2; target region 676–1859 bp; Accession No. NM_009625.2), Mm-Slc17a7-C2 (Cat. No. 416631-C2; target region 464–1415 bp; Accession No. NM_182993.2), Mm-Slc17a6-C3 (Cat. No. 319171-3; target region 1986–2998 bp; Accession No. NM_080853.3), Mm-Grid1-C1 (Cat. No. 552311; target region 396–1301 bp; Accession No. NM_008166.2), Mm-Adcyap1r1-C2 (Cat. No. 409561-C2; target region 4108–5038 bp; Accession No. NM_001025372.1), Mm-Chrna7-C3 (Cat. No. 465161-C3; target region 175–1122 bp; Accession No. NM_007390.3), Mm-Prkcd-C4 (Cat. 441791-C4; target region 334 - 1237 bp; Accession No. NM_011103.3), Mm-Sst-C4 (Cat. No. 404631-C3; target region 18–407 bp; Accession No. NM_009215.1). Following hybridization, slides were washed twice (2 min each) in wash buffer, then sequentially incubated with the amplification reagents in the HybEZ™ oven at 40 °C. Nuclei were counterstained with DAPI (20 s at RT), and slides were coverslipped and stored at 4 °C until imaging. Confocal images were acquired using a Leica Stellaris confocal microscope with a 20× objective and four channels (DAPI, Alexa Fluor 488, 550, and 647).

For duplex experiments, the following probes were used: Mm-Adcyap1 (Cat. No. 405911-C1 and 405911-C2; target region 676–1859 bp; Accession No. NM_009625.2), Mm-Slc17a7-C2 (Cat. No. 416631-C2; target region 464–1415 bp; Accession No. NM_182993.2), Mm-Slc17a6 (Cat. No. 319171-C1 and 319171-C2; target region 1986–2998 bp; Accession No. NM_080853.3), Mm-Calca-C2 (Cat. No. 578771-C2; target region 2–772 bp; Accession No. NM_007587.2), Mm-Chat (Cat. No. 408731-C1; target region 1090–1952 bp; Accession No. NM_009891.2). Following hybridization, slides were washed twice (2 min each) in wash buffer and sequentially incubated with amplification reagents. Cells were counterstained with hematoxylin, coverslipped, and examined under a bright-field microscope.

### Ex vivo whole-cell electrophysiology

Ai9-TdTomato-PKCδ mice were used for this study. Anesthesia was induced with pentobarbital, and mice were subsequently decapitated. Brains were rapidly removed and immersed in an ice-cold cutting solution containing (in mM): 92 NMDG, 20 HEPES, 25 glucose, 30 NaHCO₃, 2.5 KCl, 1.2 NaH₂PO₄, 10 MgSO₄, and 0.5 CaCl₂, saturated with 95% O₂/5% CO₂, with an osmolarity of 303–306 mOsm. Following extraction, brains were blocked and mounted in the specimen chamber of a Leica VT1200 vibratome filled with ice-cold cutting solution. Horizontal slices (300 µm thick) containing the central amygdala were sectioned. Slices were then incubated in the same cutting solution at 34°C for 5 minutes. Subsequently, slices were transferred to a chamber containing a holding aCSF solution composed of (in mM): 92 NaCl, 20 HEPES, 25 glucose, 30 NaHCO₃, 2.5 KCl, 1.2 NaH₂PO₄, 1 MgSO₄, and 2 CaCl₂ (303–306 mOsm), saturated with 95% O₂/5% CO₂, and maintained at room temperature for at least 1 hour. Slices remained in this solution until transferred to the recording chamber.

For whole-cell recordings of intrinsic excitability, glass microelectrodes (3–5 MΩ) were filled with an internal solution containing (in mM): 135 K-gluconate, 10 HEPES, 4 KCl, 4 Mg-ATP, 0.3 Na-GTP, and 0.2% biocytin. The capsular region of the central amygdala was identified under low magnification using brightfield and fluorescence illumination. Neurons for recording were selected at high magnification using differential interference contrast (DIC) optics. After achieving whole-cell access and stabilization under voltage clamp, the configuration was switched to current clamp. Series resistance (<20 MΩ) was continuously monitored by measuring the response to a -40 pA current injection. Occasionally, we observed somatic swelling or decreased tdTomato fluorescence after prolonged whole-cell recordings; however, these changes were not associated with alterations in input resistance or firing stability and were therefore not used as exclusion criteria. Cells exhibiting a >20% change in series resistance during the recording were excluded from further analysis. Recordings were obtained using a Multiclamp 700B amplifier (Molecular Devices), filtered at 2 kHz, and digitized at 20 kHz with a Digidata 1440A Digitizer (Molecular Devices).

PACAP Application to Brain Slices: to assess the intrinsic excitability of Ai9-PKCδ^+^ neurons under current clamp, a current injection protocol was applied, consisting of 1-second steps from -40 pA to 240 pA in 20 pA increments, with 7-second intervals between steps. This protocol was continuously repeated throughout the recording session. Three experimental conditions were evaluated: (1) baseline perfusion with aCSF for 15 minutes, (2) application of aCSF containing 100 nM PACAP for 15 minutes, and (3) a washout period of 15 min with standard aCSF. For each cell, five runs of the protocol in each condition were averaged. For each cell, firing current-firing frequency (I-F) curves were analyzed using a two-way repeated-measures ANOVA with treatment (baseline, PACAP, washout) and current step amplitude as within-subject factors, this criteria allowed classifying neurons as PACAP responders (n = 21) or non-responders (n = 9). A total of 30 neurons from the capsular region of the central amygdala (CeA) were recorded. After recording, the sections were drop-fixed in 4% PFA in PB for subsequent developing with streptavidin-488 and immunocharacterization.

### Transmission Electron Microscopy

Brains from C57BL wild-type mice were sectioned at 40 μm using a Leica VT1000 vibratome. The slices were rinsed in phosphate buffer (PB), incubated in 1% sodium borohydride in PB for 30 minutes to inactivate free aldehyde groups, rinsed again in PB, and then incubated in blocking solution [1% normal goat serum (NGS), 4% bovine serum albumin (BSA) in PB supplemented with 0.02% saponin] for 30 minutes. Sections were then incubated with the following primary antibodies: mouse anti-PKCδ^+^ (1:1000, BD Transduction, Cat# 610397); rabbit anti-PACAP (1:50, BMA, Cat# T4473); guinea pig anti-VAChT (1:500, Nittobo Medical, Cat# VAChT-Gp-Af1120); guinea pig anti-VGluT1 (1:500, Nittobo Medical, Cat# VGluT1-GP-Af570); rabbit anti-Dsred (1:1000, Clontech, Cat# 632496); guinea pig anti-VGluT2 (1:500, Nittobo Medical, Cat# VGluT2-GP-Af810); rabbit anti-GluD1C (1:500, Nittobo Medical, Cat# GluD1C-Rb-Af1390); rabbit anti-VGluT2 (1:500, Nittobo Medical, Cat# VGluT2-Rb-Af720-1); and rabbit anti-VAChT (1:500, Nittobo Medical, Cat# VAChT-Rb-Af670). All primary antibodies were diluted in 1% NGS and 4% BSA in PB supplemented with 0.02% saponin, and sections were incubated for 24 hours at 4°C. After rinsing in PB, sections were incubated overnight at 4°C with the corresponding secondary antibodies: Nanogold-Fab’ anti-mouse (1:100, Nanoprobes, Cat# 2002-1ML); Nanogold-Fab’ anti-rabbit (1:100, Nanoprobes, Cat# 2004-1ML); Nanogold-Fab’ anti-guinea pig (1:100, Nanoprobes, Cat# 2055-ML); and the Vectastain ABC-HRP kits for mouse, rabbit, and guinea pig IgGs (Vector Laboratories, Cat# PK-4002, PK-4001, PK-4007, respectively). Sections were rinsed in PB, then incubated in avidin-biotinylated horseradish peroxidase (HRP) complex in PB for 2 hours at room temperature. After another rinse in PB, sections were post-fixed with 1.5% glutaraldehyde for 10 minutes at room temperature, rinsed in PB, and rinsed in distilled water (to avoid introducing metals to the subsequent steps). Silver enhancement of the gold particles was performed using the HQ Silver Enhancement Kit (Nanoprobes, Cat# 2012) for 7 minutes at room temperature, followed by a PB rinse. Peroxidase activity was then visualized using 0.025% 3,3’-diaminobenzidine (DAB) and 0.003% H₂O₂ in PB for 5–10 minutes. Sections were rinsed with PB, post-fixed in 0.5% osmium tetroxide in PB for 25 minutes, washed in PB and double-distilled water, then stained with 1% uranyl acetate (freshly prepared) for 60 minutes. Dehydration was performed through a graded ethanol series and propylene oxide, followed by flat embedding in Durcupan ACM epoxy resin (Electron Microscopy Sciences, Cat# 14040,Hatfield, PA). Resin blocks were polymerized at 60°C for 2 days. Ultrathin sections (60 nm) were cut from the tissue surface using an Ultramicrotome UC7 (Leica Biosystems, Deerfield, IL) with a DiATOME® diamond knife (Diatome, Hatfield, PA). Sections were collected on formvar-coated single-slot grids and counterstained with Reynolds lead citrate. Imaging was conducted on a Tecnai G2 12 transmission electron microscope. (FEI Company, Hillsboro, OR) equipped with a OneView digital camera (Gatan, Pleasanton, CA).

### Ultrastructural analysis

Serial ultrathin sections of the capsular region of the central amygdala (CeC) were analyzed. Synaptic contacts were classified according to their morphology and immunolabel and photographed at a magnification of 6,800-13,000x. In the serial sections, a terminal containing more than 5 immunogold particles was considered as immunopositive. Pictures were adjusted to match contrast and brightness by using Adobe Photoshop (Adobe Systems).

### FIB-SEM Methodology

To assess the three-dimensional anatomical characteristics of the Calyx-like contacts, some slices were processed for electron microscopy using PACAP immunohistochemistry, slices were embedded in Durcupan resin, platinum-covered and positioned in a microscope sample holder. The electron image stack was collected using a ZEISS Crossbeam 550 FIB-SEM (Zeiss Group, Oberkochen, Germany), equipped with a high-performance field emission electron column and a gallium-ion beam column. The ion beam was accelerated at 30 kV for both rough and fine milling. A coarse trench was milled around the protective platinum (Pt) layer using a 15 nA ion current. To prevent curtain artifacts, a thin (∼1 μm thick, covering an area of 10 × 10 μm) platinum protective layer was deposited on top of the catalytic layer by injecting a platinum precursor via the gas injection system, using the ion beam at 30 kV and 3 nA.

A chevron-patterned FIB milling registration mark was milled into the protective platinum pad, and an additional FIB-mediated carbon pad was deposited to create a fiducial mark. This allowed for precise control of the slice thickness and ensured accurate 3D alignment of the image stack, resulting in distortion-free reconstruction. These processes were controlled using ZEISS ATLAS 3D Nanotomography software.

The images were acquired using an Energy-selective Backscattered (EsB) detector with the EsB grid set to 1.5 kV. The SEM beam operated at an accelerating voltage of 3 kV and a current of 1.5 nA, which was optimized for sequential removal of thin slices during the nanotomography process. The total depth of the trench created by this procedure was 100 μm. Backscattered electron images were captured in the X-Y plane, while the sequential removal of slices occurred along the Z-axis. For tomography, a pixel size of 20 nm × 10 nm was used, corresponding to a slice thickness of 20 nm.

The raw FIB-SEM dataset comprised 1568 images, which were cropped and processed into a single volume image. The final volume image was reconstructed and analyzed and segmented using ORS Dragonfly software.

### Freeze-fracture replica immunolabeling (FRIL)

Anesthetized mice were perfused transcardially using a peristaltic pump at a flow rate of 5 ml/min with 25 mM phosphate buffered saline solution (PBS) for 1 min, followed by ice cold 1% paraformaldehyde (PFA) and 15% saturated picric acid in 0.1 M phosphate buffer (PB) for 7 min. Coronal slices (130 µm thick) were cut using a vibrating microslicer (VT1000, Leica, Vienna, Austria) in 0.1 M PB. The regions containing the CeC and BNST were trimmed from the slices and immersed in graded glycerol (10%–30% in 0.1MPB) at 4°C overnight and frozen by a high pressure freezing machine (HPM 010; BAL-TEC, Balzers, Liechtenstein). Frozen samples were fractured and replicated as previously described (Schonherr, Seewald et al. 2016). The replicas were washed three times in 50 mM Tris-buffered saline (TBS, pH 7.4) containing 0.05% bovine serum albumin (BSA), 0.1% Tween-20, and 0.05% sodium azide and blocked with 5% BSA in washing buffer for 1 h at room temperature. The primary antibodies used for this study were: guinea pig polyclonal IgG raised against the 717-754 amino acid residues common to all AMPAR subunits (diluted 1:200, Frontier Science Co. Ltd, Hokkaido, Japan, cat. no. panAMPAR-GP-Af580-1) and rabbit polyclonal IgG raised against the green fluorescent protein (GFP) from the jellyfish Aequorea victoria (diluted 1:1,000, Molecular Probes-Invitrogen, cat. no. A11122, Lot. no. 1356608). Antigen-antibody complexes were identified using secondary antibodies against the species of the first antibody and conjugated to gold particles of different size: goat anti-guinea pig IgG conjugated with 5 nm gold particles and goat anti-rabbit IgG conjugated with 10 nm gold particles (both diluted 1:30, British Biocell International, Cardiff, UK). Incubation was carried out overnight at 15°C. The labelled replicas were examined using a transmission electron microscope (CM-120; Philips).

**SI Table 2.** Antibodies used in this study.

| Target | Host Species | Dilution | Source | Catalog Number |
| --- | --- | --- | --- | --- |
| **For electron microscopy** |  |  |  |  |
| DsRed | Rabbit | 1:1000 | Clontech | 632496 |
| GluD1C | Rabbit | 1:500 | Nittobo Medical | GluD1C-Rb-Af1390 |
| Green fluorescent protein | Rabbit | 1:300 | Molecular probes | A11122 |
| PKCδ | Mouse | 1:1000 | BD Transduction | 610397 |
| PACAP | Rabbit | 1:50 | BMA | T4473 |
| AMPA receptor (GluR1-4) | Guinea pig | 1:200 | Frontier Science | Pan-AMPA-GP-Af580-1 |
| VACHT | Guinea pig | 1:500 | Nittobo Medical | VACHT-Gp-Af1120 |
| VACHT | Rabbit | 1:500 | Nittobo Medical | VACHT-Rb-Af670 |
| VACHT | Rabbit | n/a | Provided by Dr. Lee Eiden (NIMH, NIH) | n/a |
| VGluT1 | Guinea pig | 1:500 | Nittobo Medical | VGluT1-GP-Af570 |
| VGluT2 | Guinea pig | 1:500 | Nittobo Medical | VGluT2-GP-Af810 |
| VGluT2 | Rabbit | 1:500 | Nittobo Medical | VGluT2-Rb-Af720-1 |
| **For light microscopy** |  |  |  |  |
| Green fluorescent protein | Rabbit | 1:1000 | Molecular probes | A11122 |
| PACAP | Rabbit | 1:2000 | BMA | T-4473 |
| VGluT1 | Guinea pig | 1:5000 | Chemicon | AB-5905 |
| VGluT2 | Mouse | 1:1000 | Chemicon | MAB-5504 |
| VGluT2 | Guinea pig | 1:2000 | Synaptic Systems | 135 404 |
| CGRP | Rabbit | 1:1000 | Phoenix Pharmaceuticals | H-015-09 |
| Neurotensin | Mouse | 1:1000 | Sigma | SAB-4200703 |
| Calretinin | Goat | 1:2000 | Swant | CG1 |
| VGluT2 | Guinea pig | 1:3000 | Synaptic Systems | AB 2301731 |

**Secondary antibodies**
| Reagent | Target | Dilution | Source | Catalog Number |
| --- | --- | --- | --- | --- |
| Nanogold-Fab' anti-mouse | Mouse | 1:100 | Nanoprobes | 2002-1ML |
| Nanogold-Fab' anti-rabbit | Rabbit | 1:100 | Nanoprobes | 2004-1ML |
| Nanogold-Fab' anti-guinea pig | Guinea pig | 1:100 | Nanoprobes | 2055-ML |
| Biotinylated Goat anti-rabbit | Rabbit | 1:500 | Vector Laboratories | BA-1000 |
| DyLight 405 Goat anti-rabbit | Rabbit | 1:500 | Jackson ImmunoResearch | 111-475-003 |
| Alexa Fluor 488 Donkey anti-goat | Goat | 1:500 | ThermoFisher | A-11055 |
| Alexa Fluor 488-Donkey anti-rabbi | Rabbit | 1:1000 | *ThermoFisher* | A-3279 |
| Alexa Fluor 594<br>goat anti mouse | mouse | 1:500 | Jackson<br>ImmunoResearch | 115-585-003 |
| Cy3 Donkey anti-guinea pig | Guinea pig | 1:500 | Jackson<br>ImmunoResearch | 706-165-148 |
| 15 nm gold anti-rabbit | Goat | 1:30 | BB International | 7190 |
| 15 nm gold anti-guinea pig | Goat | 1:30 | BB International | 1175 |

